# Oral Bacterial Translocation Promotes Lung Dysbiosis and Infection Risk after Spinal Cord Injury

**DOI:** 10.64898/2026.10.09.758070

**Authors:** Katherine A. Mifflin, Kristina A. Kigerl, Pilar Manrique, Ashley Cyr, Ajay Kumar Medipally, Zhen Guan, Kymberly M. Gowdy, Phillip G. Popovich

## Abstract

Spinal cord injury (SCI) causes systemic pathophysiology that extends beyond the injured spinal cord. Respiratory infections remain a leading cause of death after SCI and can impair neurological recovery, yet the microbial source of these infections remains poorly defined. This study defined the onset and origin of pulmonary dysbiosis after SCI by testing whether commensal bacteria move to the lung from the gut or oral cavity. In a murine model, bacterial culture, 16S rRNA sequencing and source-tracking analyses showed that SCI rapidly increased lung bacterial burden and shifted the pulmonary microbiome toward an oral-like community. Gut-associated bacteria did not measurably reach the lung through lymphatic or vascular routes. Systemic antibiotics commonly used to treat infection failed to reliably reduce post-SCI lung bacterial burden. In contrast, oral and lung-directed antibiotic delivery reduced bacteria linked to post-injury dysbiosis more effectively. These findings identify the oral cavity as a proximal source of lung dysbiosis after SCI, and support route-specific strategies to prevent respiratory infection after SCI.

**One Sentence Summary:** SCI promotes oral, but not gut, bacterial translocation to the lung, revealing a targetable source of post-injury pulmonary dysbiosis.

## INTRODUCTION

Respiratory complications following spinal cord injury (SCI) increase mortality, delay recovery, and reduce quality of life (1–3). Pulmonary infections are a leading cause of death after both acute and chronic SCI, and pneumonia is a primary reason for rehospitalization (1). These outcomes have changed little over the past 50 years, indicating that current clinical practices do not adequately prevent pulmonary complications. A lack of preventive or targeted therapies, combined with an incomplete understanding of post-injury pulmonary immunity, continues to limit progress.

Although pneumonia diagnosis after SCI can be ambiguous and treatment may be delayed, prophylactic antibiotics are rarely used because of concerns about promoting antibiotic resistance (4). This creates a clinical dilemma: patients with SCI are highly vulnerable to pulmonary infection, yet the available first-line defenses are constrained by diagnostic uncertainty, rising antibiotic resistance, and evidence that antibiotics may be less effective after CNS injury (5, 6).

Prior work shows that SCI disrupts systemic and pulmonary immune defenses, increasing susceptibility to infection, including pneumonia (7–11). In murine models of SCI, injury reduces pulmonary leukocyte numbers, suppresses transcriptional programs required for host defense, and increases lung pathogen burden (10). Reducing environmental pathogen exposure may lower infection risk in immunocompromised patients with SCI, but this strategy is unlikely to fully prevent post-injury pulmonary infections for two reasons. First, although SCI reduces pulmonary leukocyte numbers and suppresses immune responses, increasing immune cell recruitment to the lung does not prevent the post-SCI increase in pulmonary bacterial burden (10). Second, potential pulmonary pathogens, including Streptococcus pneumoniae, can reside within the normal lung microbiome (12). Thus, SCI-associated pulmonary infections may arise not only from environmental exposure, but also from dysregulation of resident commensal bacteria, in addition to intrinsic changes in the lung microenvironment.

Although the lung was once considered sterile, sequencing-based analyses have identified a distinct lung microbiome that differs from gut and oropharyngeal microbial communities. Furthermore, lung dysbiosis occurs in several pulmonary disorders (see (13) for a review) and after CNS disorders For example, after stroke, pulmonary dysbiosis occurs due to bacterial translocation from the gut (14). Whether similar changes are seen after SCI is unknown. After SCI, reduced pulmonary immune surveillance could allow lung dysbiosis to develop through expansion of resident bacteria, aspiration or translocation of oral cavity bacteria, or through dissemination of gut bacteria through lymphatic or vascular routes. Several injury-associated factors could support cross-organ bacterial transfer: SCI causes gut dysbiosis, and acute care practices such as ventilatory support, nil per os status (restricted oral intake), while reduced oral hygiene may alter the oral microbiome (15, 16). Defining the microbial source of post-SCI lung dysbiosis could guide prophylactic and targeted therapies to prevent pulmonary infection after injury.

This study tested how commensal bacteria contribute to pulmonary infection risk after SCI and identified the route by which these bacteria reach the lung. SCI caused lung dysbiosis by altering endogenous commensal communities rather than introducing new environmental bacteria. Bacteria translocated primarily from a dysbiotic oral cavity to the lung, not from the gut, distinguishing SCI from other CNS injuries in which gut-derived bacteria contribute to pulmonary dysbiosis (14). Targeted treatment of the oral cavity and lung also reduced post-SCI bacterial burden more effectively than systemic antibiotics. Together, these findings show that SCI promotes oral-to-lung bacterial transfer and identify route-specific strategies to manage pulmonary infections after SCI.

## RESULTS

### SCI rapidly remodels the pulmonary microbiome independently of environmental exposure

16S rRNA sequencing of whole lung homogenate revealed compositional remodeling of the pulmonary microbiome after SCI. By 3 days post-injury (dpi), the pulmonary microbiome of SCI mice separated from that of naïve and laminectomy (Lam) control mice, with injury explaining ∼45% of the between-group variance (**Fig. 1A**, PERMANOVA). Dispersion did not differ among groups (Fig. 1A). Analysis of between-group Bray-Curtis dissimilarity showed an overall effect of injury without significant pairwise differences, consistent with a moderate shift in beta-diversity (**Fig. 1B**). Alpha-diversity, measured by the Shannon Diversity Index, showed a non-significant downward trend after SCI, consistent with early loss of community homeostasis as the lung microbiome shifts toward enrichment of select taxa (**Fig. 1C**). Phylum-level profiling supported these findings: although relative bacterial abundance differed across conditions, specifically an increase in *Bacillota* after SCI, the same bacterial phyla were detected in each group (**Fig. 1D**).

**Figure 1.**
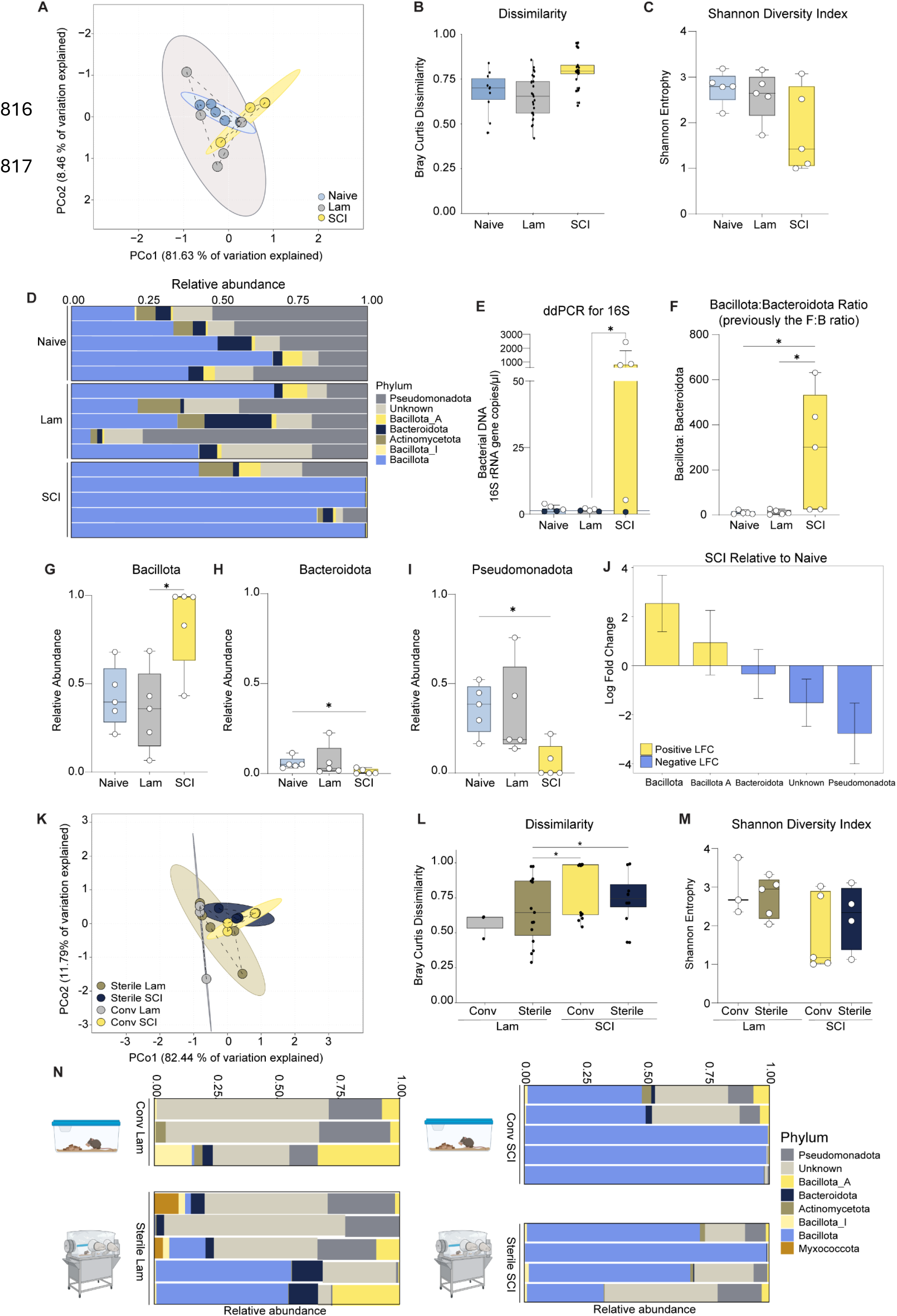
The lung microbiome changes after SCI. Whole lungs from Naive, T1 laminectomy (Lam), and T1Tx (SCI) mice were taken at 3d post-injury for 16S sequencing (n=5/group). (**A**) Principal coordinate analysis (PCoA) of Bray-Curtis distances shows SCI lungs differ from controls (PERMANOVA with 999 Permutations F_2,14_= 4.88, R^2^= 0.45, P=0.018). Each dot represents one animal plotted in the ordination space. A dispersion analysis on Bray-Curtis distances found no difference (PERMDISP F_2,12_= 1.7593, P= 0.26) Coloured circles indicate the SD for each group. (**B**) Box plots of Bray-Curtis dissimilarities. Each black dot represents one pairwise comparison. (PERMANOVA with 999 permutations, F_3,15_= 2.547397 P=0.014, pairwise comparison test with FDR Naive vs SCI q= 0.066; Lam vs SCI q = 0.066). (**C**) Box plots of Shannon Diversity Index. No significant change. Each white dot represents one animal. (H(2) = 3.62, P=0.1636). (**D**) Stacked bar graphs of lung microbes at the phylum level. (**E**) 16S rRNA gene copies per sample as measured by digital droplet PCR. Line represents the number of copies in the no-template control. Black dot samples did not pass this threshold and were not included in analyses. (H(2) = 7.000, P=0.0062, Lam vs. SCI adj. p = 0.0349) (**F**) Box plots of the ratio of all *Bacillota* to *Bacteroidota* (H(2) = 7.940, P= 0.0111, Naive vs SCI adj. p= 0.0486, Lam vs SCI adj. p=0.0400). (**G-I**) The relative abundance of *Bacillota* (**G,** H(2) = 7.340, P=0.016, Lam vs. SCI adj. p = 0.04), *Bacteroidota* (**H,** H(2) = 6.020, P=0.0431, Naive vs. SCI adj. p = 0.0486), and *Pseudomonadota* (**I,** H(2) = 6.860, P=0.0236, Naive vs. SCI adj. p = 0.04). (**J**) Log fold change (LFC) of each phylum in SCI mice compared to Naive, as per ANCOM-BC2 analysis. Changes in *Bacillota* and *Pseudomonadota* trended towards significance (p = 0.2, Holm FDR adj. p = 0.13). (**K**) PCoA of Bray-Curtis distances shows the microbial community of SCI lungs differ from Lam lungs regardless of housing in conventional (Conv) cages (n= 3 Lam (2 samples excluded due to poor sequencing, n=5 SCI) or a sterile isolators (n=5 Lam, n=4 SCI, PERMANOVA with 999 Permutations main effect injury F_1,13_= 12.2380, R^2^= 0.45, P=0.003). Each dot represents one animal. Coloured circles indicate the SD for each group. A dispersion analysis on Bray-Curtis distances found no difference (PERMDISP F_3,13_= 1.2566, P= 0.324). (**L**) Box plots of Bray-Curtis dissimilarity distances. Each data point represents one pairwise comparison. (PERMANOVA with 999 permutations, F= 3.854532 P=0.001, Pairwise comparisons FDR * q < 0.05.) (**M**) Box plots showing Shannon Diversity Index. No significant change (H(2) = 3.10, P=0.3768). Each white dot represents one animal. (**N**) Stacked bar graphs of lung phyla. Cartoon icons indicate housing conditions. *(C,E-I,M) Kruskal-Wallis with Dunn’s multiple comparison tests, * P ≤ 0.05*.

Digital droplet PCR, performed on the same samples used for sequencing in Fig. 1A, showed that SCI lungs contained the highest number of 16S rRNA gene copies, consistent with culture-based evidence of increased bacterial burden after SCI (**Fig. 1E**) (11). Additional analyses showed that SCI increased the lung *Bacillota-to-Bacteroidota* phyla ratio (B:B ratio), a metric historically described as the *Firmicutes-to-Bacteroidetes* ratio under NCBI taxonomy (**Fig. 1F**). This nomenclature differs from prior reports because this study used the Genome Taxonomy Database (GTDB) which uses genome data to classify taxonomy rather than other characteristics used previously when genomic data wasn’t available instead of NCBI, Greengenes, or SILVA taxonomies (see methods for details). Altered B:B ratios have been associated with multiple disease states (17–20). SCI increased the lung B:B ratio by increasing *Bacillota* (former *Firmicutes*) abundance (**Fig. 1G**) and reducing *Bacteroidota* abundance (**Fig. 1H**). SCI also reduced the relative abundance of Pseudomonadota (formerly *Proteobacteria*) (**Fig. 1I**). ANCOM-BC analysis corroborated these taxonomic shifts, identifying *Bacillota* as the phylum with the largest positive log-fold change and *Pseudomonadota* as the phylum with the lowest log-fold change after SCI (**Fig. 1J**). Together, these data show that SCI induces lung dysbiosis and expansion of pulmonary bacterial biomass, with increased *Bacillota* abundance driving the shift from a healthy *Pseudomonadota*-dominated community toward a dysbiotic state (17).

To test whether environmental bacteria drive SCI-induced pulmonary dysbiosis, conventionally raised mice were injured and then immediately housed either under conventional conditions or in sterile germ-free isolators. If dysbiosis reflects remodeling of resident commensal communities rather than acquisition of environmental bacteria, then SCI should disrupt the pulmonary microbiota even under germ-free housing conditions, where the only bacteria in the environment would be commensals. Consistent with this prediction, SCI induced pulmonary dysbiosis regardless of housing condition (**Fig. 1K**). As in the initial cohort, injury explained ∼45% of the between-group variance, whereas housing had no detectable effect by PERMANOVA. SCI also shifted microbial community composition relative to Lam controls (**Fig. 1L**) without changing alpha-diversity (**Fig. 1M**). Phylum-level profiling again showed increased *Bacillota* after SCI, independent of housing condition (**Fig. 1N**). Together, these data indicate that post-SCI pulmonary dysbiosis reflects remodeling of resident commensal bacterial communities rather than acquisition of new environmental bacteria.

### SCI weakens gut-associated lymphoid tissue immunity

Gut-to-lung bacterial translocation could promote pulmonary dysbiosis after SCI. Bacterial translocation promotes lung injury and infection after sepsis and stroke (14, 18, 19). SCI-induced gut dysbiosis and increased intestinal wall permeability may create conditions that favor bacterial dissemination (20, 21). This could allow intestinal bacteria or bacterial products to enter the circulation and reshape the lung microbiome after SCI. However, sustained translocation would require failure of intrinsic mucosal defense mechanisms that normally restrict bacterial escape, including immune surveillance by gut-associated lymphoid tissues (GALT), particularly immune cells in Peyer’s patches (PPs) and mesenteric lymph nodes (MLNs).

To test this, immune cells in the MLN and PPs were examined. SCI weakened GALT immunity across both major intestinal immune compartments. Analysis of PPs showed selective T lymphocyte loss, including reductions in CD4+ and CD8+ subsets, without changes in B lymphocytes, cell-type proportions, cell death, or fecal IgA (**Supp. Fig 1, Supp. Fig. 2**). Broader immune depletion was evident in MLNs, with fewer B lymphocytes and CD3+ T lymphocytes, reduced CD4+ T cell proportions, and lower tissue weight, again without increased cell death (**Supp. Fig 1, Supp. Fig. 3**). Together, these data indicate that SCI produces site-specific but coordinated suppression of GALT immune surveillance, with modest disruption in PPs and more extensive weakening of the MLN firewall that normally limits commensal bacterial dissemination.

### Gut-derived bacteria do not reach the lung through lymphatic or vascular routes after SCI

Impaired GALT immunity could permit intestinal bacteria to escape MLN containment and enter lymphatic drainage pathways that connect the gut to the lung. After bypassing MLN filtering, bacteria could move through intestinal lymphatics, drain into the thoracic duct, enter the subclavian vein, and then to the lung (**Fig. 2A**). To test this route directly, the thoracic duct was ligated before injury. If gut-derived bacteria reached the lung through thoracic duct-dependent lymphatic trafficking, ligation should reduce post-injury pulmonary bacterial accumulation (**Fig. 2A-B**). Evans blue dye injected into the hind paw confirmed successful ligation in 11/12 mice, and the mouse with incomplete ligation was excluded. By 3dpi, SCI increased lung bacterial burden relative to controls, but thoracic duct ligation did not reduce this increase (**Fig. 2C**). MLN cultures also showed no upstream bacterial accumulation after ligation (**Fig. 2D**). To determine whether lymphatic translocation might occur later, mesenteric collecting lymph fluid was gathered at 28dpi (**Fig. 2E-G**), when gut dysbiosis, pulmonary immune dysfunction, and lung bacterial burden are well established SCI (10, 20). Although SCI mice showed increased lung bacterial burden at this chronic time point, aerobic and anaerobic cultures detected almost no bacteria in mesenteric collecting lymph (**Fig. 2H-I**). Together, these findings argue against lymphatic gut-to-lung bacterial translocation as a major driver of pulmonary bacterial accumulation after SCI.

**Figure 2.**
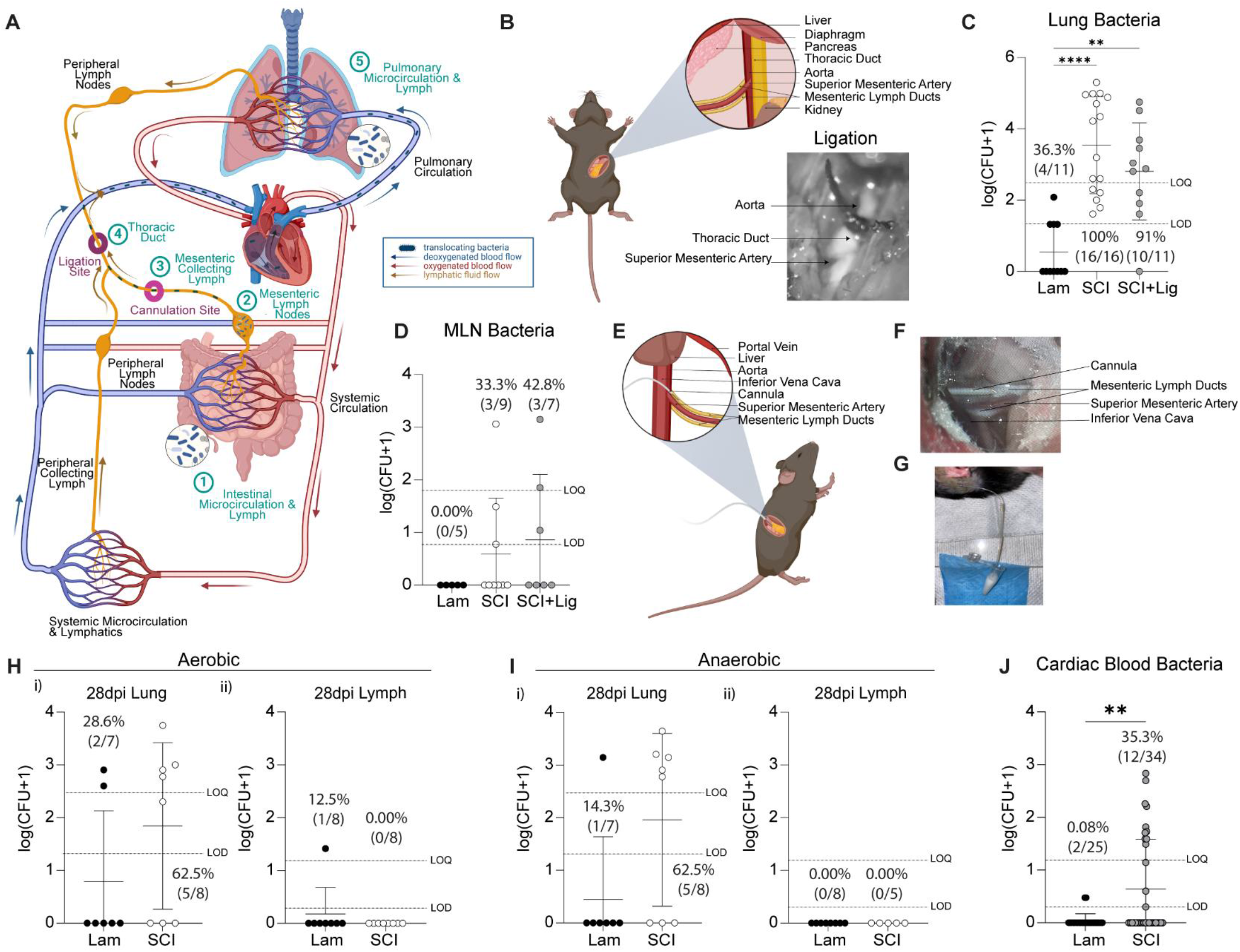
Bacteria do not translocate along the lymphatic gut-lung axis after SCI. (**A**) Schematic depicting the proposed path of translocating bacteria highlighting sites of experimental thoracic duct ligation and mesenteric lymph collection. (**B**) Schematic and grayscale surgical image of the thoracic duct ligation site. (**C**) Lung CFU quantification at 3 dpi showed ligation had no effect despite a significant increase in bacterial CFUs and proportion of plates with bacterial growth after injury (H(2) = 20.37, P <0.0001,Lam vs SCI p<0.0001; Lam vs SCI + Ligation p=0.0047, Fisher’s p=0.0002). (**D**) MLN bacterial burden did not differ. (**I**) Schematic and surgical images of the exact site of mesenteric collecting lymph cannulation in mice with T3 transection or laminectomy at 28dpi **(E-G**) Moderate increase in lung bacterial burden at 28dpi (**Hi, Ii**), but no difference in bacterial burden in the collected lymph in either aerobic (**Hii**) or anaerobic (**Iii**) blood agar cultures. (**J**) Cardiac blood showed increased bacterial burden after SCI (U = 285.5, p=0.0041, Fisher’s p = 0.0279). *Bars represent mean ± SD. Mann-Whitney U analysis(H-J). Kruskal-Wallis Test with Dunn’s multiple comparisons (C,D). The percentages above each dataset represent the proportion of plates with at least 1 CFU (growth/no growth ratio). Fisher’s Exact Two-Sided Test for proportions in (C-D,H-J). *p<0.05, ** p<0.01, **** p<0.0001. Schematic made with BioRender.com. Original surgical images. Dotted lines represent the limit of detection (LOD) and limit of quantification (LOQ) for each tissue. (C)Ligation experiments data from 2 cohorts Lam n=11, SCI n=16, SCI+Lig n=11, only one cohort in (D). (H-J) Cannulation experiments data pooled from 3 cohorts Lam n=7, SCI n=8. Data in J combined from mice used in experiments in Fig 2G and Fig 7A*.

Increased bacterial burden was observed in cardiac puncture blood (**Fig. 2J**), raising the possibility that intestinal bacteria seed the lung via a vascular gut-lung axis. In this route, bacteria would move from the intestinal circulation into the portal vein, pass through the liver and inferior vena cava, and into the pulmonary circulation (**Fig. 3A**). To test this vascular route, portal vein and inferior vena cava blood were collected sequentially at 3 dpi, along with lung and liver homogenates for culture. The inferior vena cava was ligated at the level of the kidneys so that the collected blood reflected gut-to-liver circulation rather than bacterial input from the lower body; this approach was adapted from established vascular sampling methods (22, 23). SCI increased pulmonary bacterial burden as expected, but portal vein bacteria were detected in only one mouse (**Fig. 3B-C**). Liver bacterial burden was moderately increased after acute SCI, while inferior vena cava cultures yielded minimal bacterial growth (**Fig. 3D-E**). These negative cultures could reflect limited assay sensitivity, since culture-based colony-forming unit (CFU) quantification detects only viable bacteria that grow under the selected conditions, even though it remains the standard approach for measuring live bacterial burden. To determine whether the absence of bacteria in SCI vascular samples reflected a technical floor effect, the same culture strategy was tested in cecal ligation and puncture (CLP) sepsis, a model with robust gut-derived bacterial translocation (19, 24–26). CLP increased bacterial burden in lung, portal vein blood, liver, inferior vena cava blood, and MLNs (**Fig. 3F-J**, representative culture plates in **Supp. Fig. 4)**, confirming that the CFU assay detected bacteria when vascular dissemination occurred. These results confirmed a lack of “floor effect” in the SCI data. Thus, the absence of detectable bacteria in the collected vascular samples after SCI argues against vascular gut-to-lung bacterial translocation as a major source of pulmonary bacterial accumulation. The increased bacterial burden detected in cardiac puncture blood and liver homogenates likely reflects spillover from the lung into blood returning from the pulmonary circulation, rather than dissemination of gut-derived bacteria to the lung.

**Figure 3.**
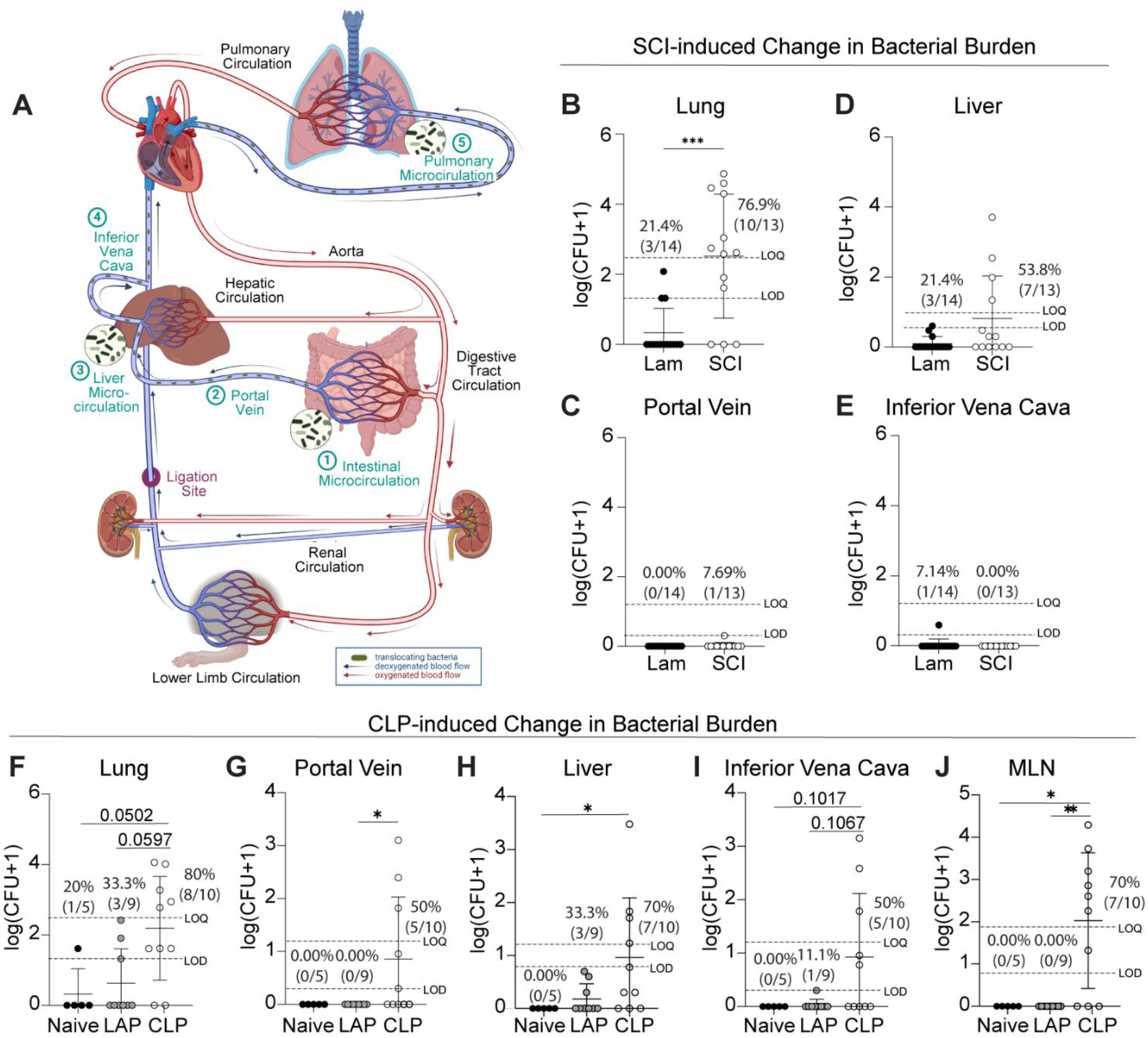
Bacteria do not translocate along the circulatory gut-lung axis after SCI. (**A**) Schematic depicts the proposed path of bacterial translocation from the gut to the lung via systemic circulation and highlights experimental tissue collection sites. (**B**) Quantification of lung CFUs (1:10 dilutions) from Lam and SCI at 3 dpi. Lung bacterial burden increased in SCI mice (U = 27.5, p=0.0005, Fisher’s p = 0.007). (**C**) Portal vein blood, (**D**) liver homogenate, and (**E**) inferior vena cava blood CFU quantification showed no difference between groups. (**F**) Lung CFU quantification from Naive, laparotomy (LAP), and mice with a cecal ligation and puncture (CLP) induced sepsis at 1 day post-infection showed a greater bacterial burden in CLP mice (H(2) = 7.979, P = 0.0185, Naive vs CLP p=0.0597; LAP vs CLP p=0.0502, Fisher’s p = 0.0725). (**G**) Bacterial burden increased in CLP mice in portal vein blood (H(2) = 8.328, P = 0.0155, LAP vs CLP p=0.02079, Fisher’s p = 0.0112), (**H**) liver (H(2) = 7.606, P = 0.0223, Naive vs CLP p=0.0294, Fisher’s p = 0.0309), (**I**) inferior vena cava blood (H(2) = 6.382, P = 0.0411, Naive vs CLP p=0.1017; LAP vs CLP p=0.1067, Fisher’s p = 0.0285), and (**J**) mesenteric lymph node (H(2) = 12.76, P = 0.0017, Naive vs CLP p=0.0208; LAP vs CLP p=0.0039, Fisher’s p = 0.0009). *Percentages above each dataset represent the proportion of plates with at least 1 CFU (growth/no growth ratio). Fisher’s exact two-sided test (B-J), Mann-Whitney U analysis (B-E). Kruskal-Wallis test with Dunn’s multiple comparisons (F-J). Bars represent mean ± SD. Schematic made with BioRender.com. Dotted lines represent the limit of detection (LOD) and limit of quantification (LOQ) for each tissue. (B-E) Data pooled from two cohorts: n=14 Lam, n=13 SCI. (F-J) Data pooled from two cohorts: n=5 Naive, n=9 Lap, n=10 CLP. *p<0.05, ** p<0.01, **** p<0.0001*.

### SCI drives oral-to-lung microbial convergence

As thoracic duct ligation did not reduce lung bacterial burden after SCI and no bacteria were detected in circulatory routes, it is unlikely that gut-to-lung translocation causes pulmonary dysbiosis. The next question was whether another body site could serve as the bacterial source of dysbiosis. The oral cavity was a strong candidate since oral bacteria can enter the lower airways through aspiration and because oral and lung microbial communities are anatomically connected. To test this possibility, potential sources were compared directly; lung, tongue, and fecal microbiomes were sequenced from naïve, laminectomy, and SCI mice. Before injury, lung and oral microbiota were related but still distinguishable, whereas gut microbiota formed a clearly separate community (**Fig. 4A-C**). Given the large tissue differences, oral and gut microbiomes were examined independently, confirming that SCI independently alters the microbial communities in each tissue (**Supp. Fig. 5**). PERMANOVA of Bray-Curtis distances confirmed significant effects of injury, tissue, and their interaction, with tissue type alone explaining most of the group differences (**Fig. 4A**). Further examination of all tissues with an analysis of multivariate dispersion found a significant effect of tissue, and a tissue-by-injury interaction, driven by reduced dispersion (or reduced variability) in SCI oral samples (**Fig. 4A**).

**Figure 4.**
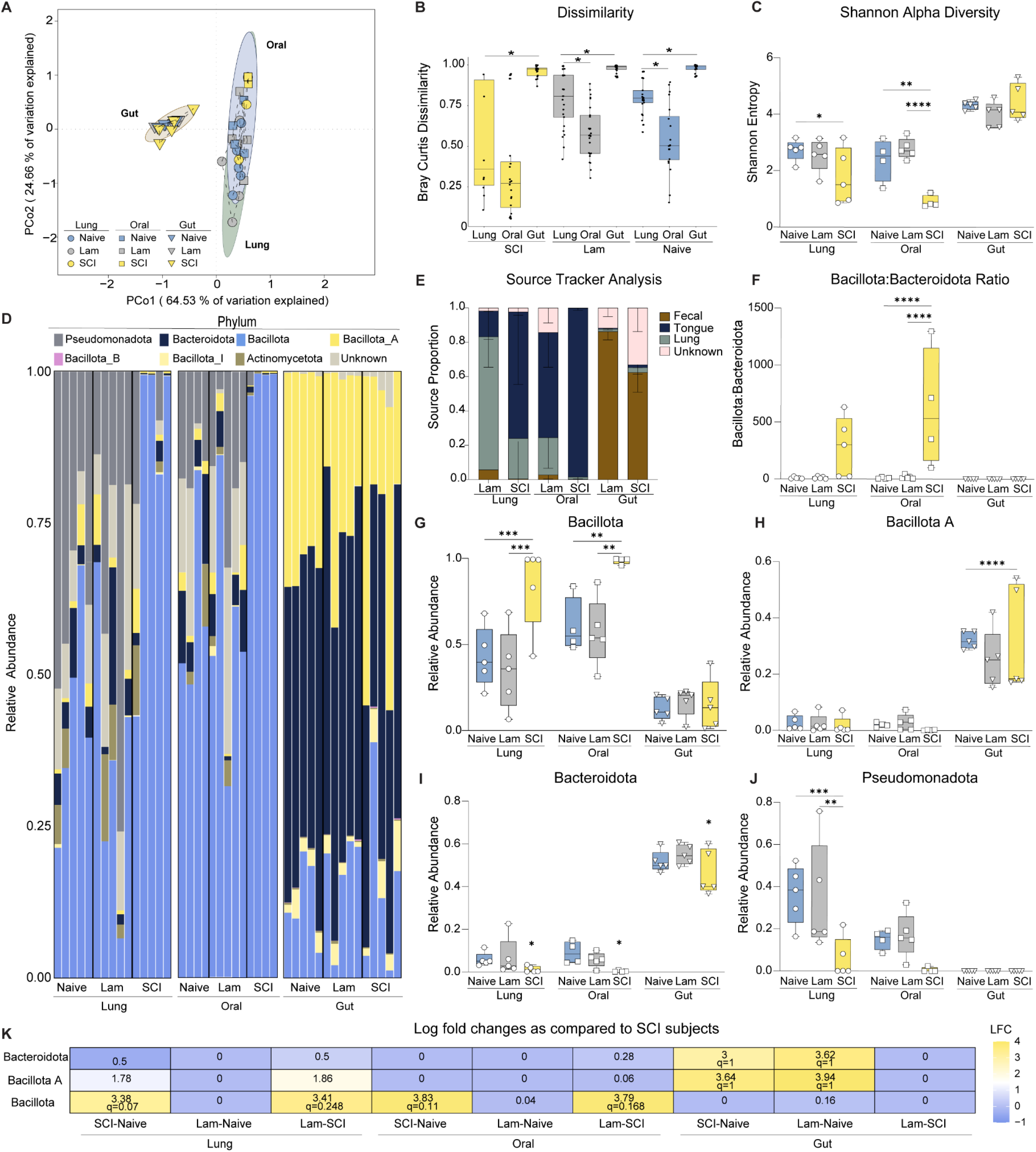
The lung microbiome mimics the oral cavity microbiome after SCI. . Whole lungs, tongues, and fecal pellets were collected from Naive, T1 laminectomy (Lam), and T1Tx (SCI) mice at 3dpi for 16S rRNA sequencing. (**A**) Principal coordinate analysis (PCoA) of Bray-Curtis distances shows separation of microbial community by injury and tissue. Community composition differed significantly by tissue (PERMANOVA with 999 permutations, R^2^ = 0.677, P =0.001), injury (R^2^ = 0.0818, P =0.001), and the tissue x injury interaction (R^2^ = 0.0617, P =0.010). Multivariate dispersion analysis detected significant effects of tissue (2-way ANOVA, F2,34 = 5.27, P = 0.010) and an tissue × injury interaction (F4,34 = 4.378, P = 0.006), but not injury alone (P = 0.0205). This was due to reduced dispersion in oral microbial communities after SCI (Tukey-adjusted P= 0.04 vs Naive, P=0.031 vs Lam). Each dot represents one animal. Outlined circles indicate the SD for each group. (**B**) Box plots showing the Bray-Curtis dissimilarities between SCI lung and all other groups. Each data point represents one pairwise comparison (PERMANOVA with 999 permutation, F= 7.139, P=0.001 with pairwise comparisons SCI lung vs gut p=0.013, SCI lung vs tongue p= 0.329, Lam lung vs oral P= 0.039, Lam lung vs gut P= 0.012, Naive lung vs tongue P = 0.026, Naive lung vs gut P=0.006). (**C**) Box plots showing Shannon Diversity Index. Each white dot represents one animal. A significant interaction of tissue x injury was seen (2-way ANOVA, F_4,34_ = 5.179, P = 0.0023). This was due to a reduction in alpha diversity in oral (Tukey-adjusted Naive vs SCI p= 0.002, Lam vs SCI P<0.0001) and lung samples (Tukey-adjusted Naive vs SCI P= 0.033). (**D**) Stacked bar graphs show microbial composition at the phylum level. (**E**) Stacked bar graphs show the relative contribution of each microbial source using Source Tracker Analysis. Error bars represent SD. (**F-J**) Box plots (**F**) the ratio of *Bacillota* to *Bacteroidota (*2-way ANOVA interaction, F_4,34_ = 4.265, P = 0.0066, Tukey’s adjusted Oral Naive vs SCI P <0.0001, Oral Lam vs SCI P <0.0001), the relative abundance of the phyla (**G**) *Bacillota* (F_4,34_ = 3.792, P = 0.0118, Lung Naive vs SCI P = 0.0009, Lung Lam vs SCI P = 0.0001, Oral Naive vs SCI P <0.0074, Oral Lam vs SCI P = 0.0021), (**H**) *Bacillota A* (main effect tissue F_2,34_ = 62.46, P <0.0001, Lung vs Gut P <0.0001, Oral vs Gut P <0.0001) (**I**) *Bacteroidota A* (main effect injury F_2,34_ = 5.687, P = 0.0074, Naive vs SCI P = 0.0215, Lam vs SCI P = 0.0124) and (**J**) *Pseudomonadota* (F_4,34_ = 2.788, P = 0.0418, Lung Naive vs SCI P = 0.0005, Lung Lam vs SCI P = 0.0012, main effect injury F_2,34_ = 8.005, P = 0.0014, Naive vs SCI P = 0.0042, Lam vs SCI P = 0.0037). (**K**) Heat map show the log fold change (LFC) of each significant phyla in SCI mice compared to Naive (ANCOM-BC2 analysis). Holms adjusted p-values (q) listed for LFC above 2. *\*P<0.05, ** P < 0.01, *** P<0.001. n=5/group except for Naive and SCI oral samples (n=4), due to exclusion for poor sequencing quality*.

Thus after SCI, lung and oral samples are no longer distinct: oral and lung communities no longer differ by Bray-Curtis dissimilarity (**Fig. 4B**) and SCI reduced Shannon diversity (**Fig. 4C**). This was most prominent in oral samples and consistent with expansion of dominant *Bacillota* taxa in both the lung and oral cavity (**Fig. 4D**). As an independent computational test of microbial origin, SourceTracker was used to estimate how much of each lung community could be attributed to lung, tongue, or fecal source communities (27). This analysis strengthened the evidence for oral-to-lung convergence: lung communities in laminectomy controls were composed mainly of lung-derived sources (∼77%), whereas SCI lung communities were composed mainly of tongue-derived sources (∼74%) and contained far fewer lung-derived bacteria (∼23%) (**Fig. 4E**).

The taxa that changed after SCI pointed to the same conclusion. Oral-lung convergence was marked by expansion of *Bacillota* (**Fig. 4D, F-K**). SCI increased the *Bacillota-to-Bacteroidota* ratio in the oral cavity and produced a similar trend in the lung (**Fig. 4F**). *Bacillota*, the dominant gram-positive phylum (dominated by the *Bacilli* class after SCI) in oral and lung samples, increased significantly in both tissues after SCI (**Fig. 4G**). By contrast, gut samples were dominated by *Bacillota A* (predominantly of the *Clostridia* class), a distinct predominantly gram-positive phylum that did not change after injury (**Fig. 4H**). SCI also reduced *Bacteroidota* across tissues (**Fig. 4I**). *Pseudomonadota*, which was enriched in oral and pulmonary microbiota but not gut communities, decreased significantly after SCI, with a tissue-by-injury interaction evident in the lung (**Fig. 4J**). Compositional differential-abundance analysis with ANCOM-BC2, which accounts for bias inherent to relative-abundance microbiome data, confirmed these compositional shifts and identified *Bacillota* as the taxon with the largest positive log fold change in both lung and oral samples (**Fig. 4K**). Overall, combined culture-based assays, 16S rRNA sequencing, and source-tracking analyses demonstrate that SCI disrupts the pulmonary microbiome through a gut-independent mechanism driven by oral-to-lung bacterial translocation. SCI-induced oral dysbiosis expands oral-associated gram-positive taxa, promoting convergence of the oral and pulmonary microbiomes after injury.

### Systemic antibiotics incompletely suppress dysbiosis-associated lung bacteria after SCI

Having identified the oral cavity as the dominant source of post-SCI pulmonary dysbiosis, the next experiments tested whether standard systemic antibiotics could prevent the spontaneous increase in lung bacterial burden after SCI. This question has direct clinical relevance because antibiotics are commonly used to manage infection risk, but the oral origin and gram-positive *Bacillota* signature of the dysbiotic lung microbiome suggested that non-targeted systemic treatment might incompletely suppress the bacteria driving oral-to-lung convergence.

Gentamicin was tested first because it is a broad-spectrum aminoglycoside often used to prevent and treat aerobic gram-negative infections (28). Daily post-injury gentamicin injections (s.c.) modestly reduced lung bacterial burden after SCI, but did not prevent pulmonary colonization; the persistence of culture-positive lungs despite treatment suggests that systemic gentamicin incompletely targets the dysbiotic lung community, potentially allowing resistant taxa to persist or expand (**Fig. 5A,B**).

**Figure 5.**
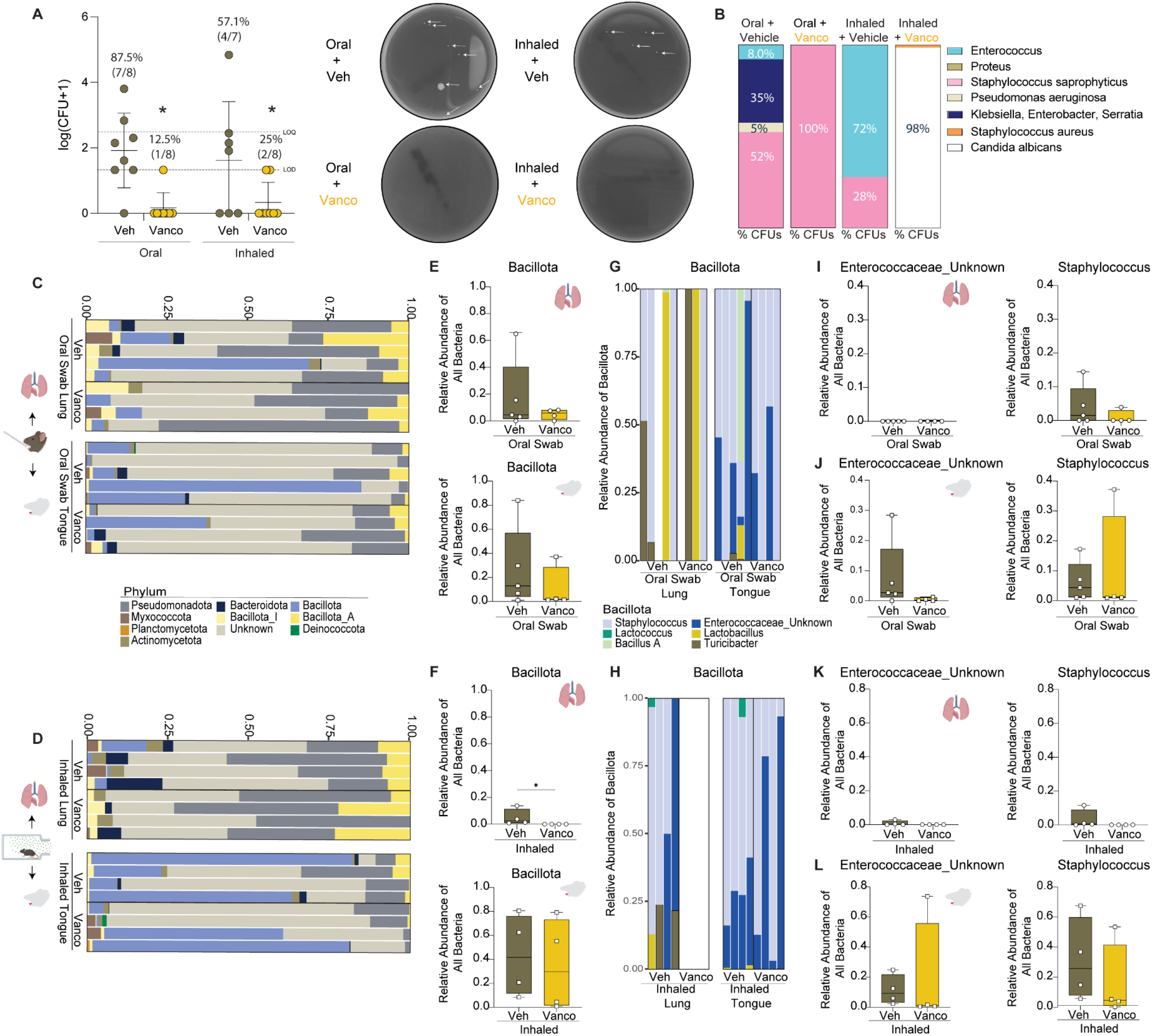
Systemic antibiotic treatment does not target dysbiosis-driving bacteria. (**A**) Lung CFUs quantification in Lam and SCI mice with and without gentamicin (Gent) or cefazolin (Cef). SCI increased lung bacterial burden (Kruskal-Wallis with Dunn’s comparisons H(3) = 11.43, P = 0.0096; Lam vs Veh P = 0.0177), and antibiotic treatment did not reduce bacterial burden. Percentages above each dataset represent the proportion of plates with at least 1 CFU (growth/no growth ratio). Fisher’s exact test of these proportions found no difference. Representative images of blood agar plates. Images were converted to greyscale to improve visualization. White arrows highlight small CFUs. (**B**) Antibiotic treatment alters bacterial composition on CHROMAgar™ plates, specifically reducing variety in the lung and increasing predominance of *Enterococcus*. Percentages represent the average proportion of each bacteria type on plates with positive cultures. (**C**) Stacked bar graphs representing the different *Bacillota* taxa composition at the genus level in lung and oral microbiome samples in SCI, Naive and Lam mice. (**D-E**) Relative abundance of all identified bacteria of the two increasing genera after SCI in the lung and oral cavity. (**D**) *Staphylococcus* (two-way ANOVA main effects of tissue, F_(1,22)_ = 10.31, P= 0.0040 and injury F(2,22) = 18.16, P< 0.0001, Tukey’s multiple comparison test lung Naive vs SCI P=0.0007, lung Lam vs SCI P=0.0026, oral Naive vs SCI P=0.0412, oral Lam vs SCI P=0.0015) and (**E**) *Enterococcaceae of unknown genus (*Two-Way ANOVA main effects of injury, F_(2,22)_ = 4.734, P= 0.00195, Tukey’s multiple comparison test Naive vs SCI P = 0.0147). Each dot represents the relative abundance in one mouse. (**F)** Vancomycin reduced lung bacterial burden in SCI mice at 3dpi (Mann Whitney U = 16, p=0.0297 on CFU). Percentages above each dataset represent the proportion of plates with at least 1 CFU (growth/no growth ratio). No proportional difference between groups. Representative images of blood agar plates. Images were converted to greyscale to improve visualization. White arrows highlight small CFUs. (**G**) Vancomycin minimally alters bacterial composition on CHROMAgar™ plates after SCI. Percentages represent the average proportion of each bacteria type on plates with positive cultures. *\*P<0.05, ** P < 0.01, *** P<0.001, n=5/group for 16S except for Naive and SCI oral samples (n=4), due to exclusion for poor sequencing quality. (A-B) Antibiotic data include two replicate experiments for each antibiotic treatment and 4 experiments total for Lam and Veh groups (n=20 Lam, n=23 Veh, n=12 Gent, n=10 Cef). (F-G) Vancomycin treatment data were pooled from two replicate experiments, n=9/group. Dotted lines represent the limit of detection (LOD) and limit of quantification (LOQ) for each tissue*.

Cefazolin (i.p.) was tested next as a second, clinically-relevant systemic antibiotic. At our university hospital, this cephalosporin antibiotic is used before spinal surgery, primarily to prevent skin infections (personal correspondence with Dr. Jan Schwab). It is also used more broadly to prevent post-surgical infections, including respiratory tract infections (29–31). Although cefazolin is not currently used as prophylaxis for pulmonary infection, prior experimental studies indicate that cephalosporins reduce pneumonia severity (30, 31). In our hands, cefazolin produced variable effects, with increased bacterial burden still evident in 50% of SCI mice (**Fig. 5A**).

Examination of lung bacterial cultures on CHROMagar™ Orientation plates revealed that gentamicin and cefazolin reduced bacterial diversity, but each failed to target specific members of *Bacillota*, especially *Enterococcus* (**Fig. 5B**). If these taxa drive pulmonary dysbiosis after SCI, this could explain why neither of these common antibiotics fully reduce or prevent lung bacterial burden in SCI mice. Further analysis of the sequencing data found that pulmonary *Bacillota* expansion was driven by a limited set of genus-level taxa. Among seven *Bacillota* taxa detected in lung and tongue samples, *Staphylococcus* and an unclassified member of *Enterococcaceae* were confirmed as the major SCI-dependent differences between control and injured mice (**Fig. 5C-E**).

Systemic (s.c.) injection of vancomycin was next tested as it is a tricyclic glycopeptide antibiotic used to treat gram-positive infections caused by *Staphylococci* and *Enterococci* (32, 33). Vancomycin reduced lung bacterial burden in ∼56% of SCI mice, but there was minimal change in the types of bacteria identified with CHROMagar™ plates (**Fig. 5F,G**). 16S rRNA sequencing of lung and tongue samples from a subset of these mice showed no overall change in the *Bacillota* phylum in lung, although treatment reduced *Bacillota* in the oral cavity (**Supp. Fig. 6A, B**). Staphylococcus abundance was unaffected by vancomycin (**Supp. Fig. 6C**), although treatment did reduce abundance of the unclassified *Enterococcaceae* taxon (**Supp. Fig. 6D**). Although vehicle-treated SCI mice showed a smaller proportion of *Bacillota* than previous cohorts, Staphylococcus and the unclassified *Enterococcaceae* taxon remained the predominant *Bacillota* taxa. Together these data demonstrated mixed effectiveness of systemic antibiotic treatment.

### Oral and airway-directed vancomycin reduces lung bacterial burden after SCI

The mixed and incomplete effects of systemic antibiotic treatment suggested that effective prophylaxis may require organ-specific interventions that directly target bacteria in the oral cavity and lung. Vancomycin, the most effective of the three antibiotics tested, was therefore administered either directly to the oral cavity as an oral rinse, or to both the oral cavity and lung by inhalation. Both approaches markedly reduced lung bacterial burden (**Fig. 6A**). In mice that retained culturable bacteria after treatment, vancomycin eliminated lung *Enterococcus* CFUs on CHROMagar™ Orientation plates in both treatment groups and *eliminated Staphylococcus saprophyticus* in the inhalation group (**Fig. 6B**). 16S rRNA sequencing of a subset of treated mice confirmed that organ-specific vancomycin administration reduced *Bacillota* abundance (**Fig. 6C-L**). This reduction reached significance only in the lung after inhalation treatment, primarily because *Bacillota* taxa were undetectable in inhalation-treated lung samples (Fig. 6F-H). Both routes of administration also eliminated detection of the unclassified *Enterococcaceae* taxon in treated lungs, consistent with the CHROMagar™ Orientation plate results (**Fig. 6B, I-L**). Together, these data demonstrate that organ-specific treatment of SCI-altered oral and pulmonary bacteria reduces lung bacterial burden more effectively than systemic antibiotic administration.

**Figure 6.**
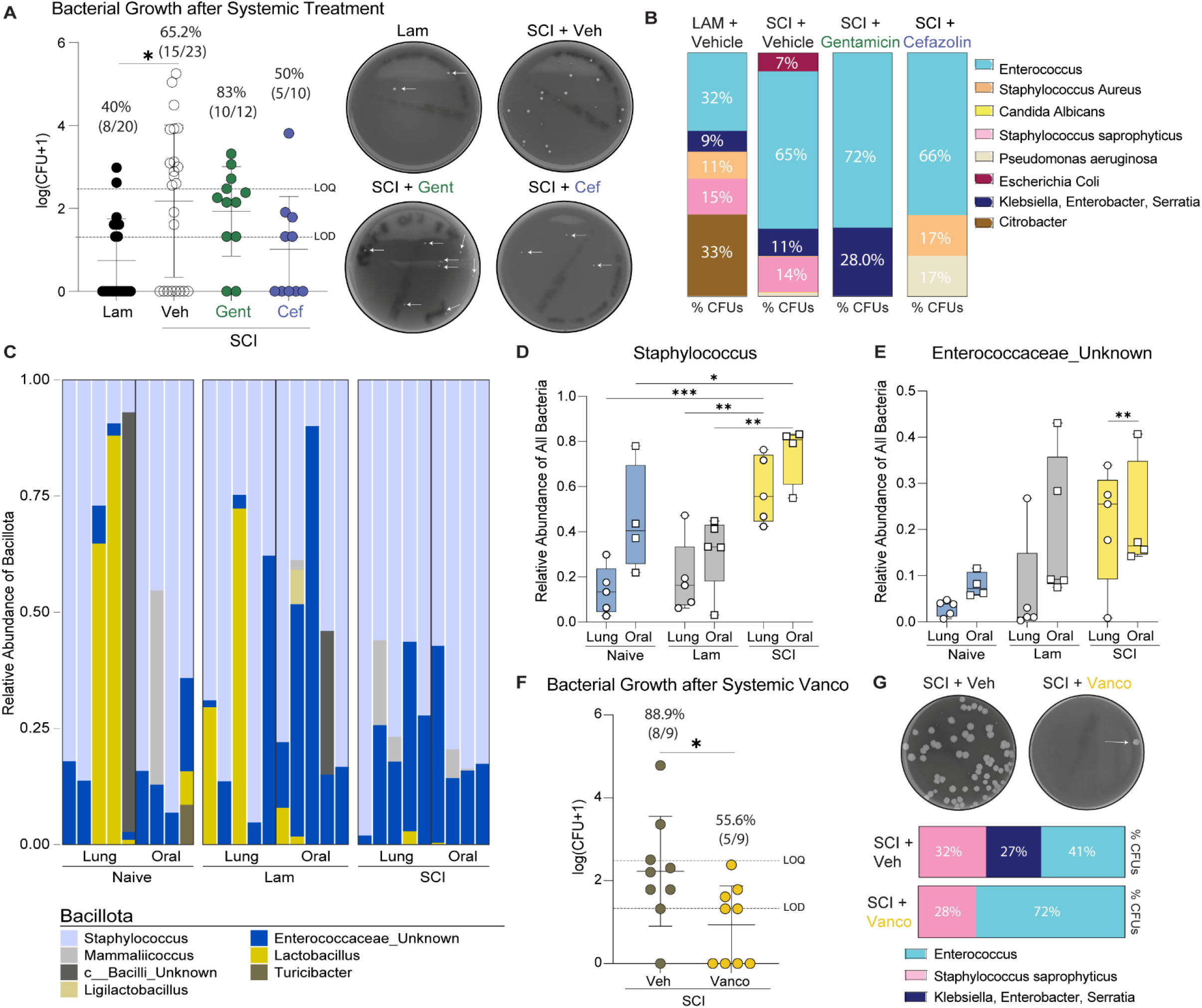
Organ-specific treatment administration reduces SCI lung bacterial burden. (**A**) Quantification of lung CFUs from SCI mice with and without vancomycin oral rinse or inhalation chamber aerosolized administration 3 dpi. Vancomycin treatment reduced lung bacterial burden regardless of route (two-way ANOVA, main effect of vancomycin, F_(1,27)_ = 14.85, P= 0.0007). Percentages above each dataset represent the proportion of plates with bacterial growth from at least 1 CFU (growth/no growth ratio). Fisher’s exact test of these proportions was significant (p = 0.0108). Representative images of blood agar plates. Images were converted to greyscale to improve visualization. White arrows highlight small CFUs. (**B**) Vancomycin reduced the *Enterococci* and most *Staphylococcus saprophyticus* on CHROMAgar™ plates. Percentages represent the average proportion of each bacteria on plates with positive cultures. (**C-D**) Stacked bar graphs represent the different phyla composition in mice treated with (**C**) oral swab vancomycin or (**D**) inhalation treatment. (**E-F**) Box plots show the relative abundance of *Bacillota* in the lung (upper) and oral cavity (lower) from (**E**) oral swab treatment or (**F**) inhalation treatment. Inhalation significantly reduced *Baillota* in lungs post-SCI (Mann Whitney U = 0, p=0.0286). (**G-H**) Stacked bar graphs represent different *Bacillota* taxa at the genus level after (**G**) oral swab or (**H**) inhalation treatment. (**I-L**) Box plots show the relative abundance of *Enterococcaceae of unknown genus* (left) *and Staphylococcus* (right) in oral swab-treated (I) lung and (J) oral cavity samples as well as inhalation-treated (**K**) lung and (**L**) oral cavity samples. Each dot represents the relative abundance in one mouse. *Antibiotic data were pooled from two replicate experiments (A&B, n=8/group for oral treatment and n=7 veh and n=8 vanco for inhaled treatment). Dotted lines represent the limit of detection (LOD) and limit of quantification (LOQ) for each tissue (A). 16S RNA Sequencing (C-H) was from a subset of mice in (A, n=4/group except n=5 for oral swab veh groups). Schematic icons made with Biorender.com, * P<0.05*

## DISCUSSION

These data identify acute oral-to-lung microbial convergence as a mechanism of pulmonary dysbiosis after SCI and a potential driver of pneumonia risk. Lung dysbiosis did not arise from environmental acquisition or gut-derived dissemination through lymphatic or vascular routes. Instead, SCI remodeled endogenous oral communities, expanded gram-positive *Bacillota* taxa, including Staphylococcus and an unclassified member of the *Enterococcaceae*, and shifted the injured lung toward an oral-like microbiome. Standard systemic antibiotics did not prevent spontaneous lung bacterial expansion, whereas oral- and lung-targeted treatments reduced dysbiosis-associated bacteria. Together, these findings place the oral cavity upstream of post-SCI pneumonia risk and support early oral- or airway-directed strategies to limit bacterial transfer to the lung.

Detecting lung dysbiosis by 3 days extends prior preclinical work showing increased pulmonary bacterial burden and infection susceptibility at acute time points after SCI (8, 10, 11). By adding 16S rRNA sequencing to culture-based assays, these data show that increased bacterial burden reflects rapid remodeling of the lung microbiome. Dysbiotic SCI lung samples also contained more 16S rRNA gene copies, indicating increased bacterial biomass. The observed increase in *Bacillota*, particularly *Staphylococcus* and an unclassified *Enterococcaceae* taxon, together with reduced *Pseudomonadota*, parallels recent reports of lung microbiome changes at 14 dpi in a T9 murine SCI model and in a clinical cohort (34). These convergent findings suggest that SCI causes early lung dysbiosis that persists beyond the acute post-injury period.

Lung dysbiosis reflected remodeling of commensal bacteria, suggesting that SCI creates a pulmonary environment prone to microbiota disruption without requiring inhaled or ingested environmental pathogens. Housing SCI mice under conventional or in sterile isolators supported this interpretation, as lung dysbiosis developed regardless of environmental exposure. Pulmonary immune suppression may help create this permissive niche, especially given the loss of pulmonary adaptive immune cells after SCI (10). For example, B lymphocytes are necessary for the antibody-dependent opsonization of gram-positive bacteria before subsequent phagocytosis, meaning their loss could promote the observed expansion of *Bacillota* taxa (35, 36). Conversely, gram-negative bacteria, like those of the *Pseudomonadota,* taxon, can be directly eliminated via the complement system without the need for opsonization, for example through the lectin complement pathway, potentially explaining their relative reduction. However, this does not fully explain why specific commensal populations expand after SCI. Within hours to days post-SCI, reduced immune surveillance may also allow low-abundance resident bacteria or aspirated oral bacteria already suited to the pulmonary environment to expand faster than newly acquired exogenous bacteria. Reduced pulmonary immune surveillance after SCI could allow low-abundance bacteria to expand and reshape the lung microbiome through interactions with other commensals. Such interactions are well described in polymicrobial lung infections in conditions like Cystic Fibrosis (37). For example, *Pseudomonas aeruginosa* and *Staphylococcus aureus* interact through complex competitive and cooperative mechanisms in cystic fibrosis airways, with *P. aeruginosa* often suppressing *S. aureus* growth (37–39). The post-SCI decrease in *Pseudomonadota*, together with expansion of *Bacillota* taxa that include *Staphylococcus*, raises the possibility that loss of competitive restraint permits different bacteria to expand in the injured lung.

These patterns of SCI-induced lung dysbiosis are reminiscent of those we have previously described in the gut after SCI and may reflect altered community structure rather than independent expansion of individual taxa (20, 21, 40, 41). As we have shown for the gut microbiome, future analysis of lung microbiome should move beyond single-taxon comparisons, especially as 16S rRNA sequencing often only allows for genera level classification. Instead, future studies should test whether SCI reorganizes pulmonary microbial networks, alters co-occurring bacterial subcommunities, or shifts encoded functions that support persistence or pathogenicity (40, 41). Genome-resolved metagenomics, metatranscriptomics, metabolomics, co-occurrence network analysis, and pathway-level modeling could identify candidate taxa that stabilize the dysbiotic lung community and reveal microbial functions that drive post-SCI pulmonary infection risk. This level of resolution could help identify oral or lung microbial species and functional changes that are selectively disrupted by SCI, rather than simply cataloging broad shifts in phyla or genera. Defining these susceptible taxa, their ecological partners, and the functions that support their persistence could reveal which members of the oral-lung microbiome drive dysbiosis and which protective commensals fail after injury. Similar precision has already enabled targeted testing of *L. johnsonii* after SCI, where restoring a gut-depleted, functionally vulnerable commensal prevented injury-induced metabolic dysfunction and systemic inflammation (40). Applied to the lung, this same strategy could guide designed probiotics, synbiotics, or niche-targeted microbial therapies that restore protective oral-lung community structure without broadly disrupting the microbiome.

Bacterial movement between organ-specific microbiomes can also drive dysbiosis. The initial hypothesis was that increased pulmonary *Bacillota* after SCI reflected bacterial translocation from the gut, since gut dysbiosis develops rapidly after SCI and includes increased *Bacillota* abundance (20, 21). These gut changes coincide with increased intestinal permeability, providing a potential route for bacteria to escape the gut and reach other organs (20). Several SCI studies report bacterial translocation after injury (34, 42), and gut-to-lung translocation occurs in sepsis, stroke, and traumatic brain injury (14, 43–47). However, many of these studies relied on detecting bacterial CFUs or endotoxin in distant organs, or on sequencing-based similarity across tissues, without systematically testing whether live bacteria occupy the anatomical routes connecting the gut and lung. To avoid this assumption, we cultured bacteria directly from blood and lymphatic pathways linking the gut to the lung after SCI, then tested further with 16S rRNA sequencing. These analyses found no evidence of gut-to-lung translocation after SCI. The culture methods were sufficiently sensitive, as the same approach detected bacteria in the vascular pathways connecting the gut and lung in a sepsis model, consistent with established gut-to-lung bacterial dissemination after sepsis (47, 48). In contrast to other neurotrauma models and prior reports in SCI, the present data show that, acutely after SCI, the oral and lung microbiomes converge, identifying the oral cavity as the more likely source of pulmonary dysbiosis. These data do however reflect reports linking oral health to respiratory diseases, including pneumonia, and to other neurological conditions like Alzheimer’s disease (49).

SCI may promote oral dysbiosis by altering the oral microenvironment, particularly saliva. Salivary protein composition, secretory IgA, flow rate, hydration, diet, and oral hygiene help shape the oral microbiome (50–52), and several of these factors can change abruptly after injury. In acute SCI care, impaired manual dexterity can limit oral hygiene, while *nil per os* status, meaning restricted oral intake, is often clinically necessary because of concerns about paralytic ileus or swallowing complications (15, 53–57). Together, these changes could disrupt salivary composition, favor expansion of dysbiosis-associated oral bacteria, and increase oral-to-lung transfer. Future studies should define how SCI changes salivary proteins, secretory IgA, flow, hydration status, and oral microbial community structure, with the goal of identifying modifiable features that reduce pulmonary infection risk. Because restricted oral intake may be medically necessary after acute SCI, oral-directed interventions, such as improved oral hygiene, hydration-focused oral care, or targeted antimicrobial rinses could be practical ways to preserve microbial balance without compromising critical care practices. Consistent with this idea, local treatment of the oral cavity and lung reduced dysbiosis-associated bacteria more effectively than systemic antibiotic delivery. Future work should determine which oral changes initiate dysbiosis, which bacteria require targeting, and how to suppress pathogenic expansion without promoting opportunistic infection.

While these data provide new insight into mechanisms that cause pulmonary dysbiosis after SCI, several limitations warrant consideration. First, all analyses were performed during the acute post-injury period. Although this interval coincides with heightened pulmonary infection risk, respiratory complications remain a lifelong consequence of SCI. Whether oral dysbiosis, pulmonary dysbiosis, and oral-to-lung bacterial transfer persist chronically is unknown. Chronic SCI is accompanied by additional physiological changes, including autonomic dysreflexia-associated immune dysfunction (8, 58–60), that could further alter pulmonary microbial communities or create alternative routes to infection. Sustained gut dysbiosis, intestinal barrier dysfunction, and immune suppression could also promote gut-to-lung bacterial translocation at later stages, as suggested by previous studies (20, 34). Second, all experiments were conducted in young adult female mice. Although no evidence suggests these results are sex-specific, validation in male animals is needed. Indeed, despite consistent evidence of SCI-induced dysbiosis in gut microbial communities, there are sex differences that could influence pathophysiology and recovery (41). Age is another important variable, as advancing age increases susceptibility to pneumonia and the average age at SCI continues to rise clinically (61, 62). Future studies should determine how sex and age influence oral-lung microbial interactions after injury. Third, this study used a severe high-level SCI model, selected to produce consistent, robust pulmonary immune dysfunction and infection susceptibility. Whether similar microbial changes occur after lower-level or incomplete SCI remains unclear. Recent evidence of pulmonary dysbiosis after T9 SCI suggests that microbiome disruption may generalize across injury severities (34), but the magnitude, timing, and mechanisms of these changes require further study.

Despite these limitations, data in this report reveal an unexpected mechanism linking SCI to pulmonary infection risk. Rather than originating in the gut, pulmonary dysbiosis arose through oral-to-lung bacterial transfer and convergence of oral and pulmonary microbial communities. This mechanistic insight had direct therapeutic implications: systemic antibiotics incompletely suppressed dysbiosis-associated bacteria, whereas oral- and lung-directed antibiotic delivery more effectively reduced pulmonary bacterial burden. Together, these data identify the oral cavity as a targetable upstream driver of post-SCI pulmonary dysbiosis and support development of localized interventions that prevent respiratory complications by interrupting bacterial transfer at its source.

## MATERIALS AND METHODS

### Study Design

The main objective of this study was to identify the source of bacterial dysbiosis of the lung after SCI. To determine whether mice develop lung dysbiosis at 3d post-injury, bacterial cultures and 16S sequencing were performed on a variety of tissues from Naïve, T1 laminectomy, and T1 complete transection mice. Mice were randomly assigned to each group using a random number generator. A cohort of these mice were also housed in conventional versus germ-free conditions to verify if lung dysbiosis develops independently of the external environment. Additionally, all mice used in 16S rRNA sequencing experiments were individuals housed. To assess the potential from gut to lung bacterial translocation after injury, additional cohorts of mice underwent thoracic duct ligation, mesenteric lymph cannulation, or cecal ligation and puncture procedures. To determine the best antibiotic treatment strategy to manage lung dysbiosis, mice with SCI also underwent treatment with multiple antibiotics (gentamicin, cefazolin, and vancomycin) administered via different routes (subcutaneously, oral swab, inhalation via aerosol exposure). For antibiotic treatment studies, mice were randomly assigned to treatment group using a random number generator. For CFU quantification and flow cytometry analysis, experiments were blinded to experimental conditions. All animal protocols were approved by the Institutional Animal Care and Use Committee of the Office of Responsible Research Practices at The Ohio State University. All experiments followed the guidelines and regulations of The Ohio State University and the National Institute of Health’s Guide for the Care and Use of Laboratory Animals.

### 16s rRNA sequencing & Analysis

Lung, tongue, and fecal samples were collected under sterile conditions and DNA extracted with DNeasy PowerSoil ProKit, followed by genomic DNA removal for lung and tongue samples (see Supplemental Methods for details). Samples were then shipped to the Environmental Sample Preparation and Sequencing Facility (ESPSF) at Argonne National Laboratory (Chicago, Illinois, USA) for sequencing.

PCR amplicon libraries targeting the V4 region of the 16S rRNA (515F-806R) were produced and DNA sequence data was then generated using Illumina paired-end sequencing.

Illumina sequencing data was then processed and analyzed using QIIME2 (v qiime2-amplicon-2024.2; (63)). In brief, samples were demultiplexed with the Golay error correction in QIIME2, then underwent sequence quality control and feature table construction using DADA2 plugin in QIIME2(64). Samples were trimmed or truncated, and sequences <250 bp were filtered out where necessary at this step. Due to the known low biomass of murine lung samples, the combined prevalence-frequency method in the QIIME2 DECONTAM plugin was applied using sequenced extraction blanks and sample DNA concentrations measured by Qubit fluorometry (65). Contaminating features were then filtered out prior to downstream analyses. Open reference clustering using the QIIME2 vsearch plugin (66) was used to cluster and assign reads to operational taxonomic units (OTUs) at 99% similarity against the Genome Taxonomy Database 220.0 (GTDB_220.0, https://gtdb.ecogenomic.org/stats/r220) reference sequences (67–70). Chimera sequences were identified using the Uchime de novo (71) and reference pipelines from vsearch (72), the QIIME2 pipeline defaults. OTUs with low abundance were filtered out (p-min frequency 100 and p-min samples 2). After chimeric and low-abundance filtering, OTUs were classified using classify-sklearn and the GTDB database version 220.0 reference as the classifier. The GTDB database was chosen as it takes a more standardized and comprehensive approach to microbial taxonomy by using genome phylogeny instead of the historical use of phenotypes to guide classification (67–70). This allowed for a more comprehensive genome-based taxonomy of culture-independent bacteria (68–70). The data was then rarefied to the sequencing depth shown in Supplemental Table 1. Sequences were then aligned, and a phylogenetic tree was created with FastTree (73) for downstream diversity analyses using the QIIME diversity core-metrics function. The SourceTracker algorithm was used to reveal microbial community origins (27). Additional statistical analyses were conducted using R (v4.2.3) and RStudio. Specifically, the Principal coordinate analysis of Bray-Curtis distances (as calculated in the QIIME2 diversity core-metrics function) was performed, and group differences assessed PERMANOVA (999 permutations) and PERMDISP. The “ggplot2” R package was used to visualize principal coordinate analyses and box plots and bar plots for these analyses. Relative abundance (ANOVA with Tukey’s multiple comparisons or Kruskal-Wallis tests with Dunn’s multiple comparisons tests, or Mann Whitney U test where appropriate) were calculated from abundance data extracted from the QIIME2 pipeline and accompanying box plot graphs were conducted in GraphPad Prism (v10) software. All relative abundances were calculated as the relative abundance to all identified bacteria, regardless of taxonomic level. For the B:B ratio analysis, the historic nomenclature of Firmicutes and Bacteroidetes was cross-referenced to the current GTDB database nomenclature (using the tools at https://gtdb.ecogenomic.org/taxon-history). As such, multiple Bacillota (Bacillota, Bacillota A, B, and I) were included in the B:B ratio Bacillota component for all analyses as they were historically considered Firmicutes phylum. The R package ANCOM-BC2 (74)was used to assess the abundance data compositionally and to determine log-fold-change of SCI compared to Naïve samples of relevant taxa.

### Statistical Analysis

General data visualization and statistical analysis were performed with GraphPad Prism (v10) software, QIIME 2-amplicon (v2024.2), and R (v4.2.3) and RStudio. Group differences were considered statistically significant at p<0.05. Sample sizes for non-sequencing experiments were determined a priori from historical data with power set to 0.8 and α = 0.05 using the program GPower (v3.1). Figures were prepared in Adobe Illustrator (v28.0), and graphical images were made in GraphPad Prism or R/RStudio and graphical schematics with Biorender.com. Two-group bacterial burdens experiments were analyzed with Mann-Whitney U analysis, while multiple-group experiments were analyzed with Kruskal-Wallis test with Dunn’s multiple comparisons where needed, while the percentage of plates with positive growth were analyzed with Fisher’s exact test. Fisher exact test was also used to analyze CHROMAgarTM bacterial compositions. Mann-Whitney U analysis was used to analyze GALT flow cytometry data, blood lymphocytes data, and Fecal IgA levels. 16S data was analyzed as described above. Data are expressed as mean ± standard deviation unless otherwise indicated. GraphPad Prism exact p-values are reported where appropriate.

## Acknowledgments

We thank L. Hudson, Y Liu, and P. Wei for technical assistance and other members of the P.G.P laboratory for their help on large experimental days. We also thank Alexandra Beauvais for her insight as our PVA spinal cord injury advocate. We thank the Genomics Shared Resource at The Ohio State University Comprehensive Cancer Center (OSU CCC), Columbus, OH, specifically their technicians P. Fadda and R. Wiemels, for conducting the digital droplet PCR analyses. Finally, we thank the technicians, specifically B. McElwain and C. Leedale, of The Ohio State University Comprehensive Cancer Center (OSU CCC). This work by the core was supported in part by the OSU CCC and the National Institutes of Health grant P30 CA16058. The content is solely the responsibility of the authors and does not necessarily represent the official views of the National Institutes of Health. We used AI to check logic and writing clarity. All data and concepts were conceived and generated by the authors.

## Funding

Craig H. Neilsen Foundation 547110 (KAM)

Paralyzed Veteran’s of America AWD-121226 (KAM)

National Institute of Neurological Disorders and Stroke – National Institute of Health R35NS111582 (PGP)

Ray W. Poppleton Endowment (PGP)

## Author contributions

Conceptualization: KAM, PGP, KAK

Data Curation: KAMs

Formal Analysis: KAM

Methodology: KAM, KAK, PM, ZG, KG, PGP

Investigation: KAM, KAK, AC, PM, ZG, AKM

Resources: KAM, KG, PGP

Software: KAM, KAK, PM

Validation: KAM

Visualization: KAM

Funding acquisition: PGP, KAM

Project administration: KAM, PGP

Supervision: PGP, KAM, KAK

Writing – original draft: KAM, PGP

Writing – review & editing: KAM, KAK, AC, PM, ZG, AKM, KG, PGP

Competing interests: The authors declare that they have no competing interests.

## Data and materials availability

### Data Availability

Source data for the animal experiments are deposited in the Open Data Commons for Spinal Cord Injury (ODC-SCI; https://odc-sci.org/) with the Accession ( dataset 1690, DOI released upon publication), and 16S raw data are uploaded to NCBI SRA under the BioProject accession number: PRJNA1539798 (under embargo until publication). All other data are available from the corresponding authors upon reasonable request.

### Code Availability

Curated data and all scripts used to analyze the data and prepare figures can be found in the following Zenodo repository: 10.5281/zenodo.23247961. (will be available publicly upon publication).

## SUPPLEMENTARY MATERIALS

Supplementary Materials and Methods

Supplementary Fig. 1 to 6

Supplementary Table S1. and S2

Supplemental References 8, 10, 20, 22, 23, 25, 26, 29, 56, 60, 75-88

