## Supplemental Materials and Methods, Supplemental Figures 1-6 for "Oral Bacterial Translocation Promotes Lung Dysbiosis and Infection Risk after Spinal Cord Injury"

### SUPPLEMENTARY MATERIALS AND METHODS

*Animal Use and Care*

Female 8-10 weeks old C57Bl/6J mice from Jackson Laboratories (Strain #: 000664) were used for all studies. Mice received commercial food pellets and chlorinated reverse osmosis water ad libitum. All animals not used for 16S rRNA sequencing were housed 5 mice/cage in ventilated microisolator cages containing corn cob bedding on a 12-hour light-dark cycle at a constant temperature (20±2°C) and humidity (50±20%). Mice used in 16S rRNA sequencing experiments were individually housed. Mice in Figure 1K-N were an expection, and were individually housed immediately post-injury in custom germ-free isolators (Park Bio) and for the reminder of the study. These germ-free housed mice received sterilized commercial food pellets (Teklad global 19% protein extruded rodent diet sterilizable; Cat # 2019S), sterilized, germ-free chlorinated reverse osmosis water ad libitum, and sterilized Inotiv Teklad Diamond soft bedding (Inotiv Cat #7089). Germ-free facilities also had a 12-hour light-dark cycle at a constant temperature of 20±2°C and humidity of 50±20%.

*Spinal Cord Injury and Post-Operative Care*

After acclimatizing for 1 week, mice underwent SCI surgery. On surgery day, intraperitoneal ketamine (80 mg/kg) and xylazine (10 mg/kg) were used to anesthetize the mice. Surgeries were completed in the morning during the light cycle. Before surgery, hair was shaved at the spinal thoracic level, followed by sequential betadine-70%-ethanol application to sterilize skin. Aseptic conditions were maintained throughout the procedure, and body temperature was supported with a warming pad. The thoracic vertebrae were exposed with a small midline incision, followed by a partial laminectomy. In T3 procedures, topical lidocaine was applied to the spinal cord to reduce spinal reflexes during the transection, and the flavum ligamentum and dura mater were opened on the spinal cord's dorsal surface to expose the transection site. Importantly, topical lidocaine was not applied in T1 transection surgeries, as we found this increases mortality. Iridectomy scissors and gentle aspiration were then used to transect the spinal cord by creating a separation between the rostral and caudal stumps of the cut cord at the T1 or T3 vertebral level (60). Transection completeness was verified under a dissection scope by the surgeon (KAM, AKM, or ZG). The muscle was sutured shut, and the skin incision was closed with surgical staples. Mice were given sterile saline (2 ml, s.q.) to aid recovery from anesthesia and placed into warm cages. All SCI mice were kept on warmers at 35.5°C to maintain body temperature for the experiment's duration. For 4d post-injury (or longer if needed), T3Tx mice received supplemental saline (1-2 ml s.q.). However, T1Tx mice required additional hydration (supplemented with 3 ml saline - s.q.: 2 ml in the morning and 1 ml in the evening). Surgical staples were removed at 7d post-injury in chronic experiments. Bladders were manually emptied twice daily throughout the experiments, and body weight and urinary pH were monitored weekly. Mice appearing dehydrated were supplemented with additional saline (1-2 ml s.q.), and pre-injury weight was maintained with both mash (mixture of water and food pellets) and high-caloric nutrient pellets as needed (only in non-16S experiments). While prophylactic antibiotics are often provided after SCI to prevent bladder infections, antibiotics could also prevent spontaneous pulmonary infections; thus, following a ULAR-approved protocol, mice were given no antibiotics unless explicitly stated. For T1 and T3 laminectomy controls (Lam), all surgical procedures were the same, but the spinal cord was not transected nor any lidocaine applied. Naïve mice underwent no surgical procedures or anesthesia.

*16s sequencing sample preparation details*

All mice used for microbiota sequencing were allowed to acclimatize to the local housing microbiome environment for two weeks prior to analysis and were individually housed to avoid cross-contamination between subjects. Fecal samples from pre-injury and at 3d post-injury were collected either directly from a scruffed mouse into a sterile tube or by collecting fecal matter into a sterile tube after placing mice in a plexiglass chamber sterilized with a chemical disinfectant (Spor-Klenz). Whole lungs and tongues were excised under sterile conditions after a terminal anesthetic overdose (2x the surgical dose) for sequencing at 3d post-injury. Expect in vancomycin antibiotic treatment experiments, where lungs were first homogenized in sterile 1xPBS for bacterial burden assessment (as described below), and then the remaining homogenate was centrifuged at 6,480g for 10 minutes at room temperature, the supernatant discarded, and the remining cell pellet immediately processed for DNA extraction as described below. Instruments were rinsed with ethanol and flame sterilized between each organ extraction and mouse. All samples were immediately flash-frozen in liquid nitrogen and stored at -80°C until DNA extraction. Prior to DNA extraction with a DNeasy PowerSoil ProKit (Qiagen #47014), whole lung and tongue samples were minced under sterile conditions with sterile surgical scissors and then bead-beaten in Qiagen PowerBead Protubes on a vortex adapter for 15 minutes. Fecal samples were also bead-beaten for 15 minutes. DNA was then extracted following the kit manufacturer's protocol. DNA blank water control samples were included in each protocol step alongside experimental samples. Given the low biomass of lung and tongue samples, we found that a secondary genomic clean-up step was necessary. This was performed using the New England Biolabs NEBNext® Microbiome DNA Enrichment Kit to facilitate host eukaryotic DNA removal. DNA concentration was then determined using a Qubit Fluorometer assay, and samples were diluted to 50ng/µl in 20µl. All samples were then shipped to the Environmental Sample Preparation and Sequencing Facility (ESPSF) at Argonne National Laboratory (Chicago, Illinois, USA) for sequencing using either the MiSeq (Illumina, data in Figures 1A-K,4, S5, 5) or NextSeq 2000 (Illumina, Figures 1 K-N, S6, 6) sequencing platforms.

Briefly, and according to Argonne National Laboratories, PCR amplicon libraries targeting the 16S rRNA encoding gene present in metagenomic DNA were produced using a barcoded primer set adapted for the Illumina HiSeq2000 and MiSeq (75). DNA sequence data was then generated using Illumina paired-end sequencing. Specifically, the V4 region of the 16S rRNA gene (515F-806R) was PCR amplified with region-specific primers that included the sequencer adapter sequences used in the Illumina flowcell (75–

79) The forward amplification primer also contained a twelve base barcode sequence that supported pooling different samples in each lane (75–79). Each 25 μL PCR reaction contained 9.5 μL of PCR (Certified DNA-Free), 12.5 μL of New England Biosystem's LongAmp® Hot Start Taq 2X Master Mix, 1 μL Golay barcode tagged Forward Primer (5 μM concentration, 200 pM final), 1 μL Reverse Primer (5 μM concentration, 200 pM final), and 1 μL of template DNA. The conditions for PCR were as follows: 94 °C for 3 minutes to denature the DNA, with 35 cycles at 94 °C for 45 s, 50 °C for 60 s, and 72 °C for 90 s; with a final extension of 10 min at 72 °C to ensure complete amplification. Amplicons were then quantified using PicoGreen (Invitrogen) and a plate reader (Infinite 200 PRO, Tecan). Once quantified, volumes of each of the products were pooled into a single tube so that each amplicon was represented in equimolar amounts. This pool was then cleaned up using AMPure XP Beads (Beckman Coulter), and quantified using a fluorometer (Qubit, Invitrogen). After quantification, the molarity of the pool was determined and diluted down to 2 nM, denatured, and diluted to a final concentration of 6.75 pM with a 10% PhiX spike for sequencing on the Illumina MiSeq. Amplicons were sequenced on a 151bp x 12bp x 151bp Nextseq run using customized sequencing primers and procedures (75).

*Digital Droplet PCR*

DNA samples were diluted to 10 ng/ul, and 2 ul was combined with 1X ddPCR™ EvaGreen Supermix (Bio-Rad Cat #186-4034) and 150 nM of each primer (F: 5'-GCAGGCCTAACACATGCAAGTC-3', R: 5'-CTGCTGCCTCCCGTAGGAGT-3') in a total reaction volume of 22 ul. Droplets were generated with 20 ul of this mix and Droplet Generation Oil for EvaGreen (Bio-Rad Cat #1864005) using the QX200 Droplet Generator. Samples were amplified under the following cycling conditions: 95 °C for 5 min; 50 cycles of 95 °C for 30 s, 60 °C for 2 min; 4 °C for 5 min; 90 °C for 5 min; 4 °C hold, with a ramp rate of 2 °C/s between all steps. Droplets were counted in a QX200 Droplet Reader, and the data was analyzed using QX Manager 2.1 software (Bio-Rad, USA).

*Mouse GALT Flow Cytometry*

After an anesthetic overdose (2x the surgical dose), the peritoneal cavity was opened, mesenteric lymph nodes (MLNs) and Peyer's Patches (PPs) were identified, excised using a dissection scope, and placed into cold RPMI media (Fisher Cat # 32404014) with 10% FBS (Fisher Cat # A5669701). The surrounding mesenteric fat was then trimmed from the MLNs under a dissection scope, and the MLNs were weighed. Tissues were then digested in Collagenase A (1mg/ml; Sigma Cat No 10103578001) and media for 15 minutes at 37°C on a rotating shaker at 30rpm, after which the reaction was quenched with 0.5M EDTA. The digested MLNs and PPs were then passed through a 40µm filter using the back of a 3ml syringe and the filter was rinsed with 1xPBS. The suspension was centrifuged (5min, 500g, 4°C), supernatant discarded, cells resuspended in 1ml flow buffer (0.1M PBS containing 1%BSA and 0.01% sodium azide) and then aliquoted into individual flow cytometry tubes for staining. Prior to staining, all cells were incubated with an Fc Blocking solution (rat anti-CD16/32 (1:10)) for 10-15 mins. Cells were then stained with a primary antibody master mix to identify immune cell populations (see Table I). After a 30-minute incubation in the dark, cells were washed with flow buffer, centrifuged (5min, 500g, 4°C), and resuspended in 1% paraformaldehyde for 20 minutes at 4°C for fixation. Cells were then washed with flow buffer, centrifuged (5min, 500g, 4°C), and resuspended in 300µl of flow buffer. Samples were passed through a 40µM filter to remove debris before analysis. Fluorosphere counting beads (50µl per sample; BD Liquid Counting Beads, NC0613836) were added for absolute cell number quantification. Appropriate unstained and isotype controls were used to determine background autofluorescence and antibody specificity, and samples with single-stained beads were used as compensation controls for each antibody. Compensation was applied to remove spectral overlap, and isotype and fluorescent minus one (FMO) controls were used to define the gating strategy (See Supplemental Figure 1). Cells were analyzed using a BD LSRFortessa flow cytometer, FACSDiva, and FlowJo software. FSC-A versus SSC-A was used to exclude debris, and doublets were excluded using FSC-H versus FSC-W plots. Dead cells were identified using the Zombie GreenTM Viability Kit per the manufacturer's instructions (BioLegend Cat # 423112) and excluded from the final analyses.

**Table SI**. Flow Cytometry Antibodies

| Antigen | Host, amount per sample, working concentration | Vendor, catalog number |
|---|---|---|
| mFC Block (Cd16/32) | Rat, 0.5µl, 0.025mg/ml | BD Biosciences, 553142 |
| mCD45-Pacific Blue | Rat, 1µl, 0.001mg/ml | Invitrogen, MCD4528 |
| mCD3-PE | Rat, 1µl, 0.002mg/ml | BD Biosciences, 555275 |
| mB220/CD45R-PE-Cy7 | Rat, 1µl,0.002mg/ml | BD Biosciences 552772 |
| mCD4-APC | Rat, 1µl,0.002mg/ml | BD Biosciences, 553051 |
| mCD8a- PerCP-Cy5.5 | Rat, 1µl,0.002mg/ml | Fisher, 45-0081-82 |

*Blood Lymphocyte Quantification*

After terminal anesthesia (as described above), blood was collected via cardiac puncture (approximately 400µl/mouse) and placed into EDTA-coated tubes. Whole blood cell counts were obtained using a HESKA element HT5TM machine, an automatic hematology analyzer that uses flow cytometer, impedance, and colorimetry to perform a 5-part white blood cell differential for multiple species, like mice.

*Fecal IgA Quantification*

Fecal samples were aseptically collected (as described above, 1-2 pellets per mouse) and immediately flash-frozen in liquid nitrogen and stored at -80°C prior to processing. Fecal samples were homogenized in 500ul sterile 1xPBS (Fisher Cat # 10010049) with a handheld homogenizer with a sterile disposable pestle. Samples were then briefly spun in a bench-top centrifuge to allow particulates to settle. The supernatant from each sample was collected and diluted 1:1000 in 1xPBS. A mouse uncoated IgA ELISA kit (Invitrogen Cat t# 88-50450) was then used per the manufacturer's protocol. Plates were read at the 450nm wavelength on a SpectraMAX190 and analyzed using SoftMax Pro Software (Molecular Devices, San Jose, CA).

*Multi-organ Bacterial Burden Assessment*

Multiple organs and blood samples were collected for bacterial burden assessment. Pulmonary bacterial burden was determined by quantifying spontaneous bacterial growth using methods outlined in previous studies (8, 10, 20). Briefly, collected lungs were homogenized into a single cell suspension in 10ml sterile 1xPBS by grinding lungs with the back of a 10ml syringe through a 100µm sterile filter. The median lobe of the liver was collected for liver assessments and homogenized in 500µl sterile 1xPBS by grinding the liver with the end of a 10ml syringe through a 70µm sterile filter. Mesenteric lymph nodes were collected and then homogenized in 5ml sterile 1xPBS by homogenizing the tissue through a 100µm filter by grinding with a 3ml syringe. All cardiac blood samples were collected from the right ventricle with 28G sterile needles, placed into uncoated sterile 1.5ml tubes, and then immediately plated to prevent clotting. Lymphatic fluid was collected as described in the relevant sections below.

All tissues were collected and homogenized under sterile conditions to prevent external bacterial contamination. Lung homogenates (500µl of the suspended solution diluted 1:10), liver homogenates (100µl of undiluted homogenate), blood samples (20µl undiluted), mesenteric lymph node homogenates

(200ul undiluted), and collected lymph (~10-20 µl  undiluted) were then plated onto blood agar (Fisher Cat #R01201) or CHROMagarTM Orientation plates (Fisher Cat #B254102) under flame sterilization and incubated under aerobic conditions at 37°C for 24 hours for blood agar plates and 48 hours for CHROMagarTM Orientation plates for aerobic bacterial growth. For anaerobic conditions, separate plates were placed into air-tight containers with a MitsubishiTM AnaeroPack-CO2 gas generation sachet to create an anaerobic environment (Thermo Scientific Cat #R681007) before being incubated as described. Blood agar plates were used for total colony forming units (CFU) counts, while CHROMagarTM plates were used to identify different bacteria. After incubation, plates were imaged, and colonies were manually counted. Total CFUs/ml were calculated based on the plate dilution factor where appropriate. The limit of detection was set to 20 CFUs/ml for lung, 5 CFUs/ml for mesenteric lymph nodes, 50 CFUs/500µl for liver, 1 CFUs/20µl for blood, and 1 CFUs/10µl for collected lymphatic fluid. Based on a calculation of limit of blank mean (LOB, based on 59 control plates across all experiments) of 0.273 CFUs and a standard deviation of 1.496, the limit of quantification (10x the LOB standard deviation) was set to 300 CFUs/ml for lung, 75 CFUs/ml for mesenteric lymph nodes, 150 CFUs/500µl for liver, 15 CFUs/20µl for blood, and 15 CFUs/10µl for collected lymphatic fluid. For each experiment, blank control plates and PBS control plates were included to identify any potential contamination during tissue preparation or plating procedures.

*Thoracic Duct Ligations*

Approximately 1 hour prior to anesthesia, mice were gavaged with sterile sesame oil (0.1ml) and given a 1 ml subcutaneous injection of sterile 0.9% saline on the right dorsal side to visualize lymph vessels during surgery. After mice received surgical anesthesia (described above), the abdomen and back were shaved and scrubbed with betadine-70%-ethanol solutions. Aseptic conditions were maintained during the surgical procedure, and a warming pad was used to maintain body temperature. Mice were placed on their right side, and an approximate 2 cm subcostal incision on the left abdomen was performed to open the abdominal cavity and expose the mesenteric lymph vessels, aorta, and thoracic duct. Mesenteric lymph vessel and the thoracic duct were identified due to their distinct white color, after prior gavage with sesame oil. The spleen, liver, pancreas, and stomach were retracted to one side, and the kidney and adrenal gland to the other, using two small retractors and saline-moistened surgical gauze. Just above the adrenal gland, a small incision was made to expose the aorta and it was freed from surrounding fatty tissue using S&T vessel-dilating forceps (both angled 10° and 45°; Fine Scientific Tools) and sterile cotton swabs. This exposed the thoracic duct at its junction with the mesenteric lymph duct and the intersection of the aorta and the superior mesenteric artery. Once the thoracic duct was gently separated from the aorta using the vessel-dilating forceps, a 6.0 silk suture thread was looped under the thoracic duct to ligate it. Successful ligations saw immediate lymphatic fluid pooling below the ligation. Thoracic duct ligation procedures were based on (80–82). Control (laparotomy) mice underwent the same surgical procedures but with the thoracic duct and surrounding vasculature being left untouched. The peritoneum was then closed with 5-0 Biosyn sutures, and the skin was closed with Michel wound clips. The mice in the ligation plus SCI groups were then flipped onto their stomachs, and the T1 SCI procedure was performed as described above. To aid recovery, mice were injected subcutaneously with pre-warmed 0.9% saline solution (0.1ml) using a 25G needle and placed on a heating pad until anesthesia recovery. At the experimental end point, mice were injected with 30µl of a 2.5% solution of Evans blue subcutaneously in the left hind paw 20 minutes prior to euthanasia to verify ligation success. Ligations were considered successful if dye was only pooled in the lymph below the ligation site, while control mice had dye throughout the thoracic duct (80).

*Mesenteric Lymph Duct Cannulations*

After receiving either, a T3 laminectomy or complete T3 transection (described above) 28 days prior, mice in the cannulation studies were subjected to mesenteric lymph cannulation. A T3 transection injury was used for the chronic timepoint to improve survival post-injury. Similar to the thoracic duct ligations described above, mice were gavaged with sterile sesame oil and given subcutaneous saline on the left dorsal side to help visualize the lymph vessels. Surgical techniques for cannulation were adapted from (83). Prior to surgery, cannulation catheters were prepared by cutting the distal tips of Alzet Osmotic Pumps rat intrathecal catheter (short, Cat # 0007741) to a bevel point to mimic a needle for insertion into the lymphatic duct and to reduce the total length to approximately 5-7 inches. After reaching the anesthetic surgical plane and being prep for surgery as described above for the ligations, mice were placed on their left side, and an ~2 cm subcostal incision on the right abdomen was performed to open the abdominal cavity and expose the peritoneal cavity. Using sterile cotton swabs, the intestines and colon were moved to the left side and held in place with a moistened sterile surgical gauze and a small retractor. This allowed for superior mesenteric artery and mesenteric lymph duct visualization. The retroperitoneal attachment to the liver was cut with surgical microscissors to better visualize the surgical field. An 18G needle was then pushed through the skin from inside the peritoneal cavity to the outside right abdominal wall, ensuring that the needle is completely parallel to the superior mesenteric artery. The previously prepared catheter was then passed through the needle into the cavity, and the 18G needle was removed to leave the catheter in place. A small incision was then made in the lymph duct using Vannes scissors. The beveled catheter was then inserted bevel-up into the mesenteric lymph duct (approximately 2-3mm) using microforceps to gently hold the lymphatic tissue. Lymph fluid would then immediately start flowing into the catheter, if not, a small amount of suction could be applied to the catheter line using a needle attached to the catheter's terminal end. The flowing lymph fluid was then collected in sterile PCR tubes (Fisher NC0926537) for approximately 2-3 hours until terminal anesthesia overdose. The collected lymph fluid was then plated for bacterial analysis as described above.

*Portal Vein and Inferior Vena Cava Blood Collection*

The techniques used to collect blood from the portal vein and the inferior vena cava are modified from (22, 23). Briefly, after terminal anesthesia, the abdominal cavity was opened along the midline up to the diaphragm. Two sterile cotton swabs were then used to move the intestines and expose both the vena cava at the level of the kidney and the portal vein. Curved forceps were used to hook under the inferior vena cava at the level of the kidney and pull through a thread piece to form a light ligation (not yet cutting off blood flow). A sterile cotton swab was then used to clear the portal vein, and a sterile 1 ml syringe with a 28G needle was gently inserted into the portal vein, parallel to the vessel, entering from the bottom direction and moving upwards towards the diaphragm. The syringe was then drawn back to collect ~20-100µl of blood from the portal vein, which was then immediately plated as above. The previously placed thread ligation on the vena cava was then tightened to cut off the blood supply from the anterior body. The ribs and diaphragm were then opened, exposing the lungs, heart, and upper portion of the inferior vena cava just below the heart. A new 1 ml syringe with a 28G needle was then inserted into the inferior vena cava to collect blood (~20-50µl) which was immediately plated. Lung and liver samples were then excised from the same animals and homogenized for plating as described above.

*Cecal Ligation and Puncture Model of Sepsis*

To ensure that bacterial translocation could be detected using our culturing techniques, we employed a cecal ligation and puncture (CLP) model of moderate sepsis as a positive control. The model was based on those described in (25, 26). Briefly, mice were prepped for surgery as described above and a ~1.5 cm vertical midline laparotomy was performed under aseptic conditions to expose the cecum with the adjoining intestine. The cecum was then tightly ligated with a 6.0 silk suture. The distance from the distal end of the cecum to the ligation point was measured to be <1 cm to ensure a moderate infection (25, 26). The cecum was then perforated twice with a 19-G needle on the same cecum side and below the ligation, approximately 1/3 down each time. The cecum was then gently squeezed to extrude a small amount of feces from the perforation sites and then returned to the peritoneal cavity. The peritoneum was then sutured closed with 5-0 Biosyn sutures, followed by suturing the skin with 5-0 Biosyn sutures. The mice were injected with pre-warmed 0.9% saline solution (0.1ml) using a 25G needle to induce the hyperdynamic phases of sepsis (25, 26). Mice were then placed on a heating pad until they recovered from anesthesia and were kept there overnight to maintain body temperature. Laparotomy mice (LAP) followed the same procedure but without the ligation and puncture. Tissue was collected 24h after the CLP procedure. Immediately prior to tissue collection, the peritoneal cavity of each mouse was lavaged with 5ml sterile betadine solution to prevent contamination of collected tissues from the septic peritoneal fluid. The peritoneal fluid was then extracted with a 21G needle, the peritoneal cavity opened, and tissues extracted and prepped as described above.

*Antibiotic Treatments*

Mice receiving our standard prophylactic antibiotic treatment were administered subcutaneous gentamicin sulfate (gentamicin, 5mg/kg; Vet One # 13985-633-04, the vehicle was 0.9% sterile saline) beginning at the time of surgery and then once daily until the study's end. Mice receiving antibiotics used as surgical prophylactic received intraperitoneal Cefazolin (250 mg/kg; Millipore Sigma Supelco PHR1291500MG, vehicle was 0.05% EtOH in sterile 0.9% saline, as drug stocks (167mg/ml) were dissolved initially in 33% ETOH then diluted to 23.8mg/ml with sterile saline prior to administration) starting 30 minutes prior to injury and then once daily for 3d post-injury (29, 84, 85). For vancomycin studies (vancomycin hydrochloride, Fisher AAJ6279006) studies, mice were either administered vancomycin subcutaneously (s.c.,180 mg/kg), orally by swabbing an antibiotic-soaked cotton swab in the oral cavity for 30 seconds (as in (56), oral swab., 180 mg/kg), or via a whole-body inhalation chamber (Hinners-style exposure chambers, 50mg in 5ml saline for 25min exposure). Dosing began at 6-8h post-injury, followed by dosing every 12 hours until the end of the experiment (86–88). Vehicle-treated mice received an equivalent volume of sterile 0.9% saline.

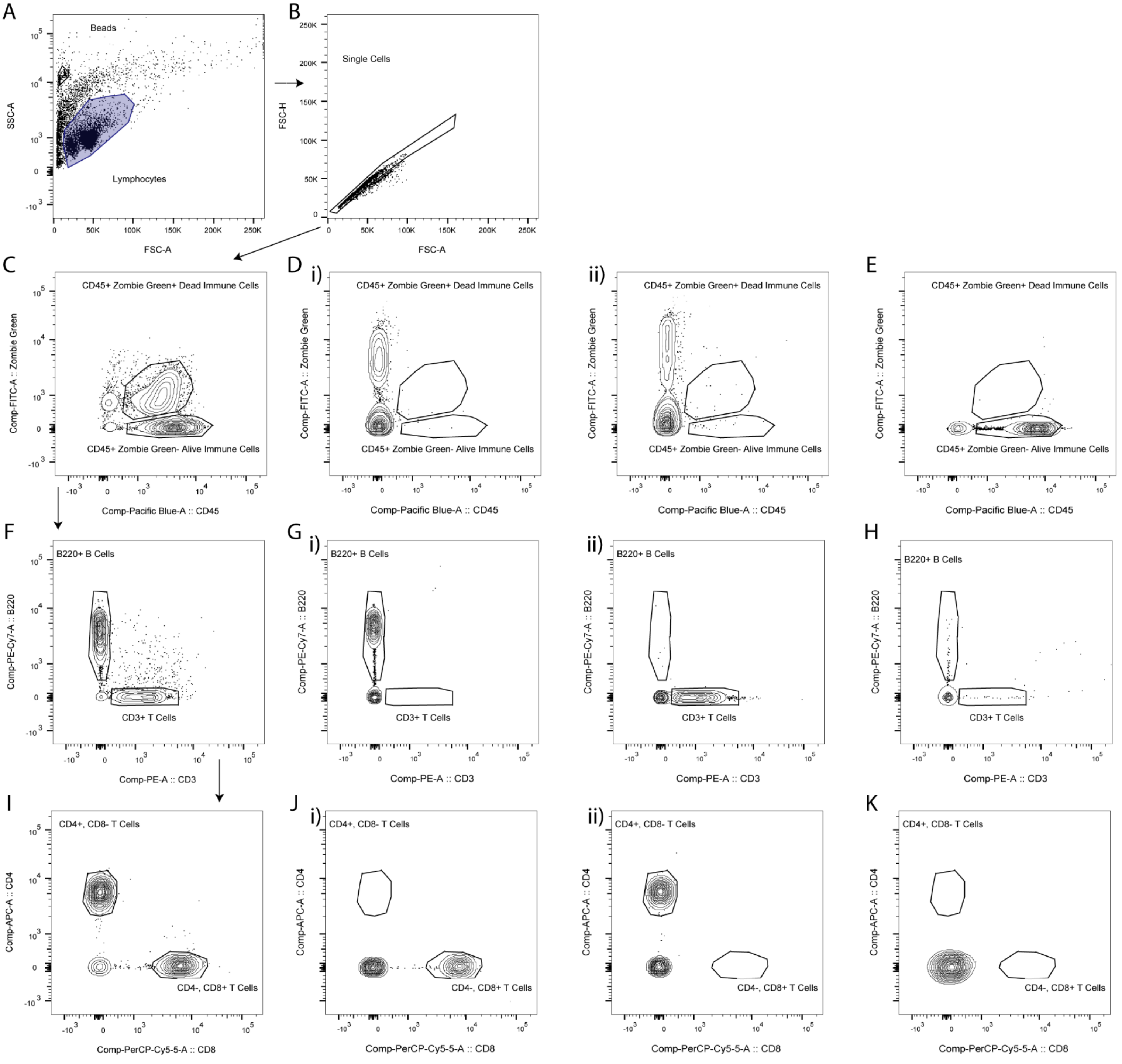
A
Beads
Lymphocytes
SSC-A
FSC-A
B
Single Cells
FSC-H
FSC-A
C
CD45+ Zombie Green+ Dead Immune Cells
CD45+ Zombie Green- Alive Immune Cells
Comp-FITC-A :: Zombie Green
Comp-Pacific Blue-A :: CD45
D i)
CD45+ Zombie Green+ Dead Immune Cells
CD45+ Zombie Green- Alive Immune Cells
Comp-FITC-A :: Zombie Green
Comp-Pacific Blue-A :: CD45
ii)
CD45+ Zombie Green+ Dead Immune Cells
CD45+ Zombie Green- Alive Immune Cells
Comp-FITC-A :: Zombie Green
Comp-Pacific Blue-A :: CD45
E
CD45+ Zombie Green+ Dead Immune Cells
CD45+ Zombie Green- Alive Immune Cells
Comp-FITC-A :: Zombie Green
Comp-Pacific Blue-A :: CD45
F
B220+ B Cells
CD3+ T Cells
Comp-PE-Cy7-A :: B220
Comp-PE-A :: CD3
G i)
B220+ B Cells
CD3+ T Cells
Comp-PE-Cy7-A :: B220
Comp-PE-A :: CD3
ii)
B220+ B Cells
CD3+ T Cells
Comp-PE-Cy7-A :: B220
Comp-PE-A :: CD3
H
B220+ B Cells
CD3+ T Cells
Comp-PE-Cy7-A :: B220
Comp-PE-A :: CD3
I
CD4+, CD8- T Cells
CD4-, CD8+ T Cells
Comp-APC-A :: CD4
Comp-PerCP-Cy5-5-A :: CD8
J i)
CD4+, CD8- T Cells
CD4-, CD8+ T Cells
Comp-APC-A :: CD4
Comp-PerCP-Cy5-5-A :: CD8
ii)
CD4+, CD8- T Cells
CD4-, CD8+ T Cells
Comp-APC-A :: CD4
Comp-PerCP-Cy5-5-A :: CD8
K
CD4+, CD8- T Cells
CD4-, CD8+ T Cells
Comp-APC-A :: CD4
Comp-PerCP-Cy5-5-A :: CD8

**Supplemental Figure 1. Flow cytometry gating strategy for Gut Associated Lymphoid Tissue (GALT).** (**A**) SSC vs FSC gating eliminates debris and separates cells and counting beads from lymphocytes. (**B**) FSC-H vs FSC-A gating eliminates doublets. (**C**) Identification of alive (Zombie Green -) immune cells ($CD45^+$) and elimination of dead cells (Zombie Green +). (**Di**) Fluorescent Minus One (FMO) control for Pacific Blue-CD45 staining. (**Dii**) FMO control for Zombie Green-FITC (Fluorescein) dye. (**E**) Isotype control for CD45+ immune cells. (**F**) Identification of $B220^+/CD45^+$ B lymphocytes and $CD3^+/CD45^+$ T lymphocytes. (**Gi**) FMO control for PE (Phycoerythrin)-CD3 antibody (**Gii**) FMO control for PE-Cy7 (Phycoerythrin-cyanine7)-B220 antibody. (**H**) Isotype control for $B220^+/CD45^+$ B lymphocytes and $CD3^+/CD45^+$ T lymphocytes. (**I**) Identification of $CD4^+/CD8^-/CD3^+/CD45^+$ and $CD4^-/CD8^+/CD3^+/CD45^+$ T lymphocytes. (**Ji**) FMO control for APC (Allophycocyanin)-CD4 antibody. **(Jii**) FMO control for PerCp-Cy5.5 (Peridinin-Chlorophyll Protein Complex- Cyanine 5.5) - CD8 antibody (**K**) Isotype control for $CD4^+/CD8^-/CD3^+/CD45^+$ and $CD4^-/CD8^+/CD3^+/CD45^+$ T Lymphocytes.

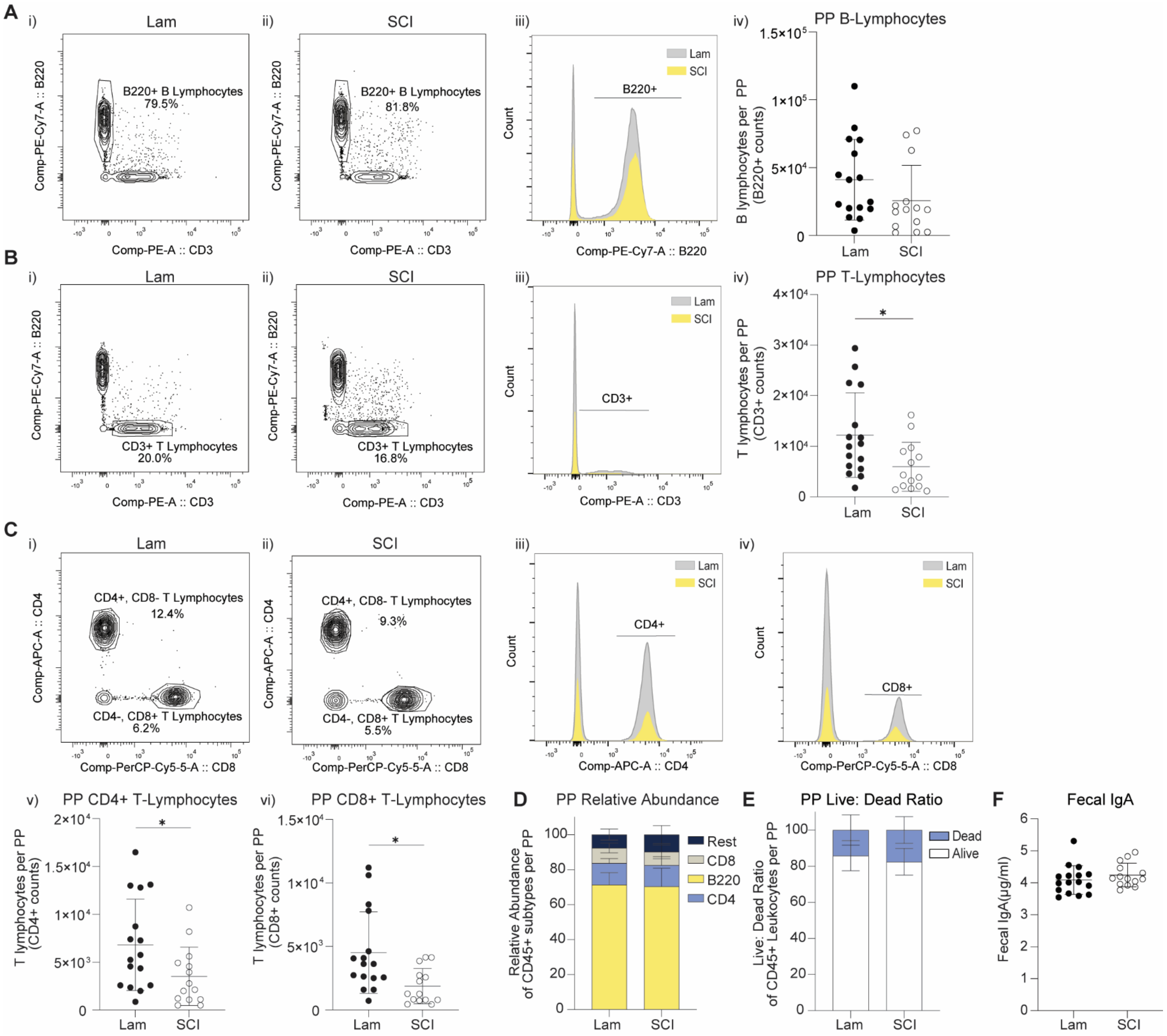
A
i) Lam
B220+ B Lymphocytes 79.5%
ii) SCI
B220+ B Lymphocytes 81.8%
Comp-PE-Cy7-A :: B220
Comp-PE-A :: CD3
iii)
Lam
SCI
B220+
Count
Comp-PE-Cy7-A :: B220
iv) PP B-Lymphocytes
B lymphocytes per PP (B220+ counts)
Lam
SCI
B
i) Lam
CD3+ T Lymphocytes 20.0%
ii) SCI
CD3+ T Lymphocytes 16.8%
iii)
CD3+
Count
Comp-PE-A :: CD3
iv) PP T-Lymphocytes
T lymphocytes per PP (CD3+ counts)
C
i) Lam
CD4+, CD8- T Lymphocytes 12.4%
CD4-, CD8+ T Lymphocytes 6.2%
ii) SCI
CD4+, CD8- T Lymphocytes 9.3%
CD4-, CD8+ T Lymphocytes 5.5%
Comp-APC-A :: CD4
Comp-PerCP-Cy5-5-A :: CD8
iii) CD4+
iv) CD8+
v) PP CD4+ T-Lymphocytes
T lymphocytes per PP (CD4+ counts)
vi) PP CD8+ T-Lymphocytes
T lymphocytes per PP (CD8+ counts)
D PP Relative Abundance
Relative Abundance of CD45+ subtypes per PP
Rest
CD8
B220
CD4
E PP Live: Dead Ratio
Live: Dead Ratio of CD45+ Leukocytes per PP
Dead
Alive
F Fecal IgA
Fecal IgA(μg/ml)

**Supplemental Figure 2. SCI reduces *Peyer's Patch* lymphocytes.** (**A**) Representative flow plots and quantification of total Peyer's Patch (PP) B220$^+$ B lymphocytes, (**B**) CD3$^+$ T lymphocytes (U = 54, p=0.0152), and (**C**) CD4$^+$ (i-iii) and CD8$^+$ (i-iv) T lymphocyte subsets. (v) Number of live CD4$^+$ (U = 61, p=0.0334) and (vi) CD8$^+$ (U = 51, p=0.0100) T lymphocytes per PP. (**D**) Relative abundance of B220$^+$ B lymphocytes, CD4$^+$, and CD8$^+$ T lymphocytes did not differ between groups. (**E**) Relative abundance of live (CD45$^+$/Zombie$^-$) and dead (CD45$^+$/Zombie$^+$) leukocytes in PPs show no difference in cell death. (**F**) Fecal IgA levels measured by ELISA did not differ. *Data were pooled from 3 cohorts ( n=16 Lam, n=14 SCI). Mann Whitney U analysis, * $P \leq 0.05$. Representative flow plots show population gating and % of total live immune cells. Bars represent mean ± SD.*

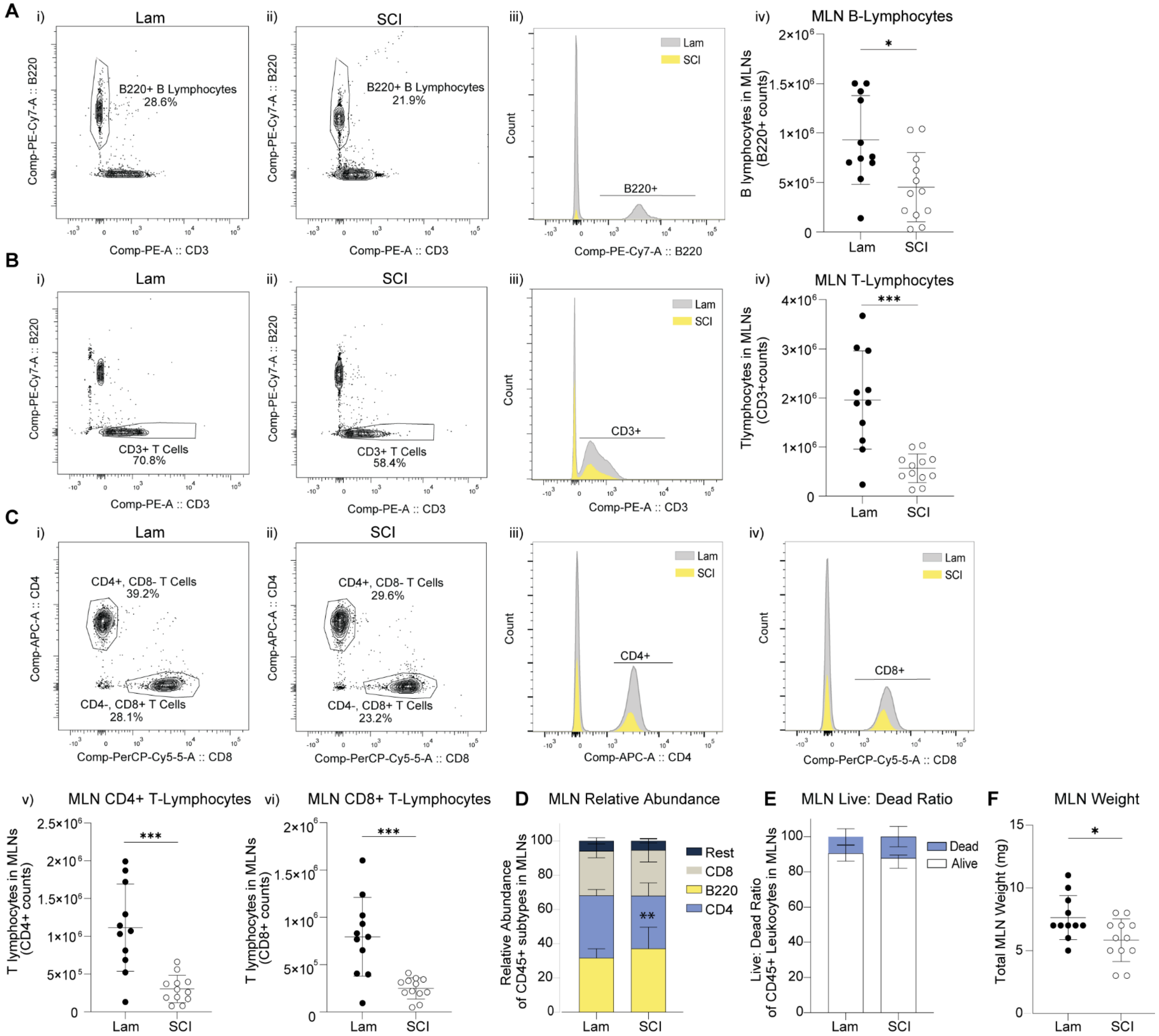
A
i) Lam
ii) SCI
iii)
iv) MLN B-Lymphocytes
B220+ B Lymphocytes 28.6%
B220+ B Lymphocytes 21.9%
Comp-PE-Cy7-A :: B220
Comp-PE-A :: CD3
B220+
Count
Comp-PE-Cy7-A :: B220
B lymphocytes in MLNs (B220+ counts)
B
MLN T-Lymphocytes
CD3+ T Cells 70.8%
CD3+ T Cells 58.4%
CD3+
Comp-PE-A :: CD3
Tlymphocytes in MLNs (CD3+counts)
C
CD4+, CD8- T Cells 39.2%
CD4-, CD8+ T Cells 28.1%
CD4+, CD8- T Cells 29.6%
CD4-, CD8+ T Cells 23.2%
Comp-APC-A :: CD4
Comp-PerCP-Cy5-5-A :: CD8
CD4+
CD8+
v) MLN CD4+ T-Lymphocytes
T lymphocytes in MLNs (CD4+ counts)
vi) MLN CD8+ T-Lymphocytes
T lymphocytes in MLNs (CD8+ counts)
D MLN Relative Abundance
Relative Abundance of CD45+ subtypes in MLNs
Rest
CD8
B220
CD4
E MLN Live: Dead Ratio
Live: Dead Ratio of CD45+ Leukocytes in MLNs
Dead
Alive
F MLN Weight
Total MLN Weight (mg)
Lam
SCI

**Supplemental Figure 3. SCI reduces Mesenteric Lymph Node lymphocytes.** (**A**) Representative flow plots and total counts of mesenteric lymph node (MLN) B220$^+$ B lymphocytes ($U = 26$, $p=0.0126$), (**B**) CD3$^+$ T lymphocytes ($U = 12$, $p=0.0004$), and (**C**) CD4$^+$ (i-iii) and CD8$^+$ (i-iv) T lymphocyte subsets. (v) Quantification of live CD4$^+$ ($U = 12$, $p=0.0004$) and (vi) CD8+ ($U = 12$, $p=0.0004$) T lymphocytes in MLNs. (**D**) Proportion of CD4$^+$ T lymphocytes decreased after SCI ($U=28$, $p=0.0188$), whereas proportions of B220$^+$ B lymphocytes and CD8$^+$ T lymphocytes did not differ. (**E**) Relative abundance of live (CD45$^+$/Zombie$^-$) and dead (CD45$^+$/Zombie$^+$) leukocytes in MLNs showed no difference in cell death. (**F**) MLN weight decreased after SCI ($U = 33$, $p=0.0373$*). Data pooled from 2 cohorts ( n=11 Lam, n=12 SCI). Mann Whitney U Analysis, * $P \leq 0.05$, ** $P < 0.01$, *** $P < 0.001$. Representative flow plots show population gating and % of total live immune cells. Bars represent mean ± SD.*

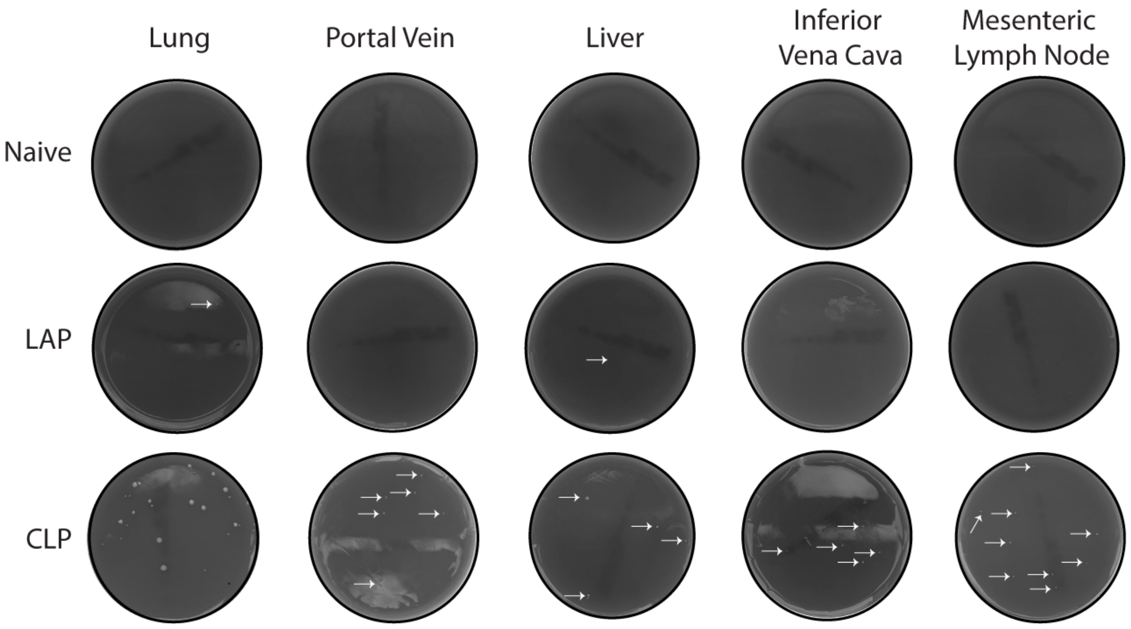


**Supplemental Figure 4. Bacterial translocation after cecal ligation and puncture.** Representative images of blood agar plates from mice modeling sepsis with a cecal ligation and puncture (CLP), control laparotomy surgery (LAP), or no surgery (Naive). Colors were converted to greyscale to improve visualization. White arrows highlight small CFUs. CLP increased bacterial burden throughout the body.

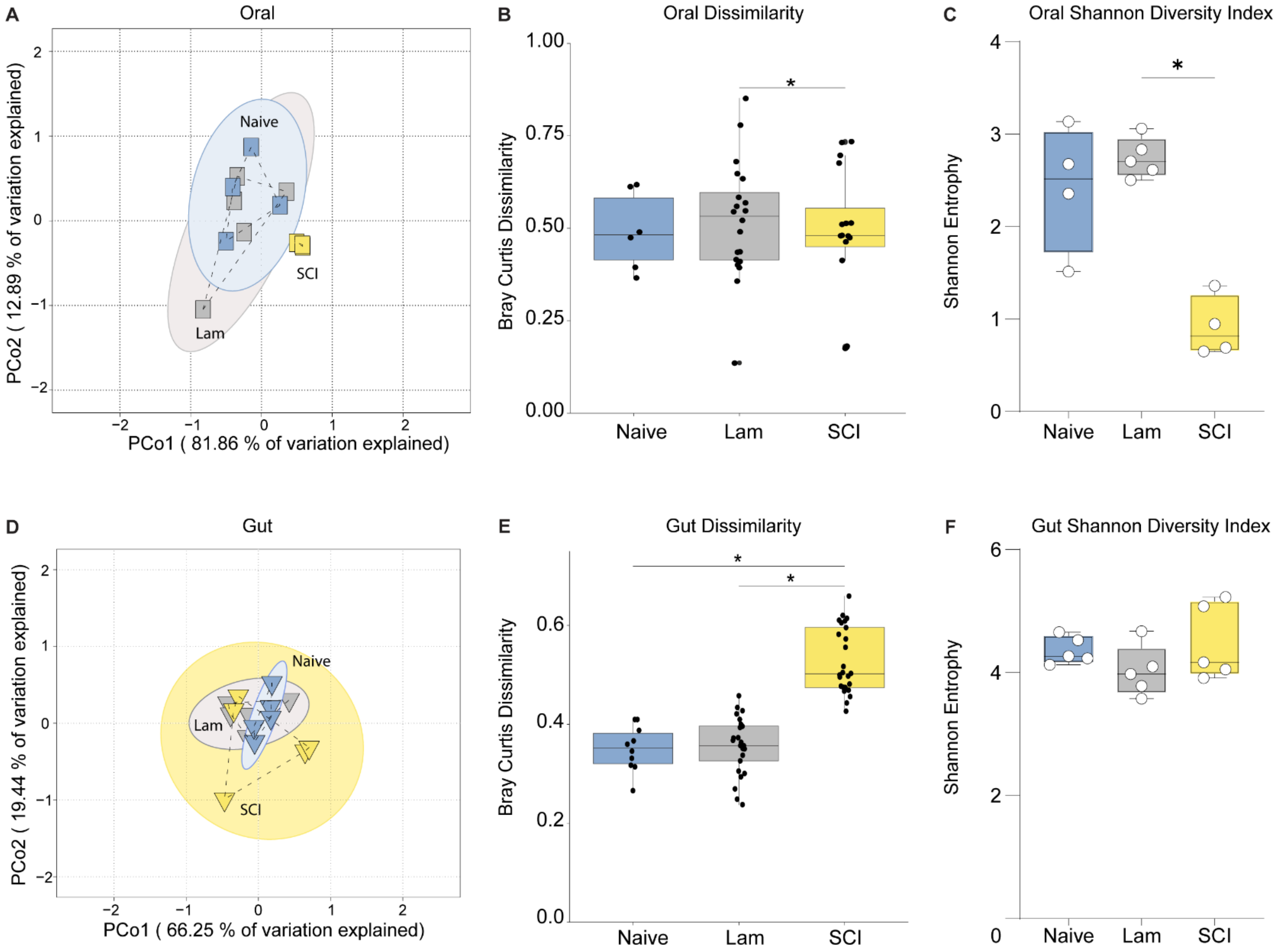
A
Oral
Naive
SCI
Lam
PCo1 ( 81.86 % of variation explained)
PCo2 ( 12.89 % of variation explained)
B
Oral Dissimilarity
Bray Curtis Dissimilarity
1.00
0.75
0.50
0.25
0.00
*
Naive
Lam
SCI
C
Oral Shannon Diversity Index
Shannon Entropy
*
Naive
Lam
SCI
D
Gut
Naive
Lam
SCI
PCo1 ( 66.25 % of variation explained)
PCo2 ( 19.44 % of variation explained)
E
Gut Dissimilarity
Bray Curtis Dissimilarity
*
*
Naive
Lam
SCI
F
Gut Shannon Diversity Index
Shannon Entropy
Naive
Lam
SCI

**Supplemental Figure 5. SCI alters the gut and oral microbiomes.** (**A**) Principal coordinate analysis (PCoA) of Bray-Curtis distances show differences in the oral microbial communities among Naive, Lam, and SCI mice (PERMANOVA with 999 permutations, $R^2 = 0.566$, P =0.007). Each dot represents one animal. Coloured circles indicate the SD for each group. Multivariate dispersion also differed among groups (PERMDISP, $F_{2,10} = 3.84$, P = 0.055), driven by a reduction in SCI dispersion (Naive vs SCI P = 0.001). (**B**) Box plots showing the Bray-Curtis dissimilarities between Naive and experimental groups (Lam and SCI) for oral cavity samples. Each data point represents one Bray-Curtis dissimilarity comparison between individual samples in each group. A higher score indicates greater diversity (PERMANOVA with 999 permutation, $F_{2,10} = 2.277$, P = 0.031 with pairwise comparisons Lam vs SCI P = 0.051). (**C**) Box plots showing Shannon Diversity Index in oral samples found reduced alpha diversity after injury (Kruskal-Wallis with Dunn's multiple comparisons H(2) = 7.981, P = 0.0068, Lam vs SCI P = 0.019). Each white dot represents one animal. (**D**) PCoA of Bray-Curtis distances shows the gut microbial communities of Naive, Lam, and SCI mice do not differ (PERMANOVA with 999 permutations, $R^2 = 0.123$, P =0.487). Each dot represents one animal. Coloured circles indicate the SD for each group. However, multivariate dispersion was significant (PERMDISP, $F_{2,12} = 12.966$, P = 0.002) and driven by increased dispersion in SCI samples (Lam vs SCI P=0.028, Naive vs SCI P = 0.002). (**E**) Box plots showing the Bray-Curtis dissimilarities between Naive and experimental groups for gut microbiome samples. Each data point represents one Bray-Curtis dissimilarity comparison between individual samples in each group. (PERMANOVA with 999 permutation, $F_{2,12} = 2.941$, P = 0.002 with pairwise comparisons Lam vs SCI P = 0.012, Naive vs SCI P= 0.012) (**F**) Box plots showing Shannon Diversity Index in gut samples found no difference in alpha diversity. Each white dot represents one animal. **P<0.05 n=5/group except for Naive and SCI oral samples (n=4), due to exclusion from poor sequencing quality.*

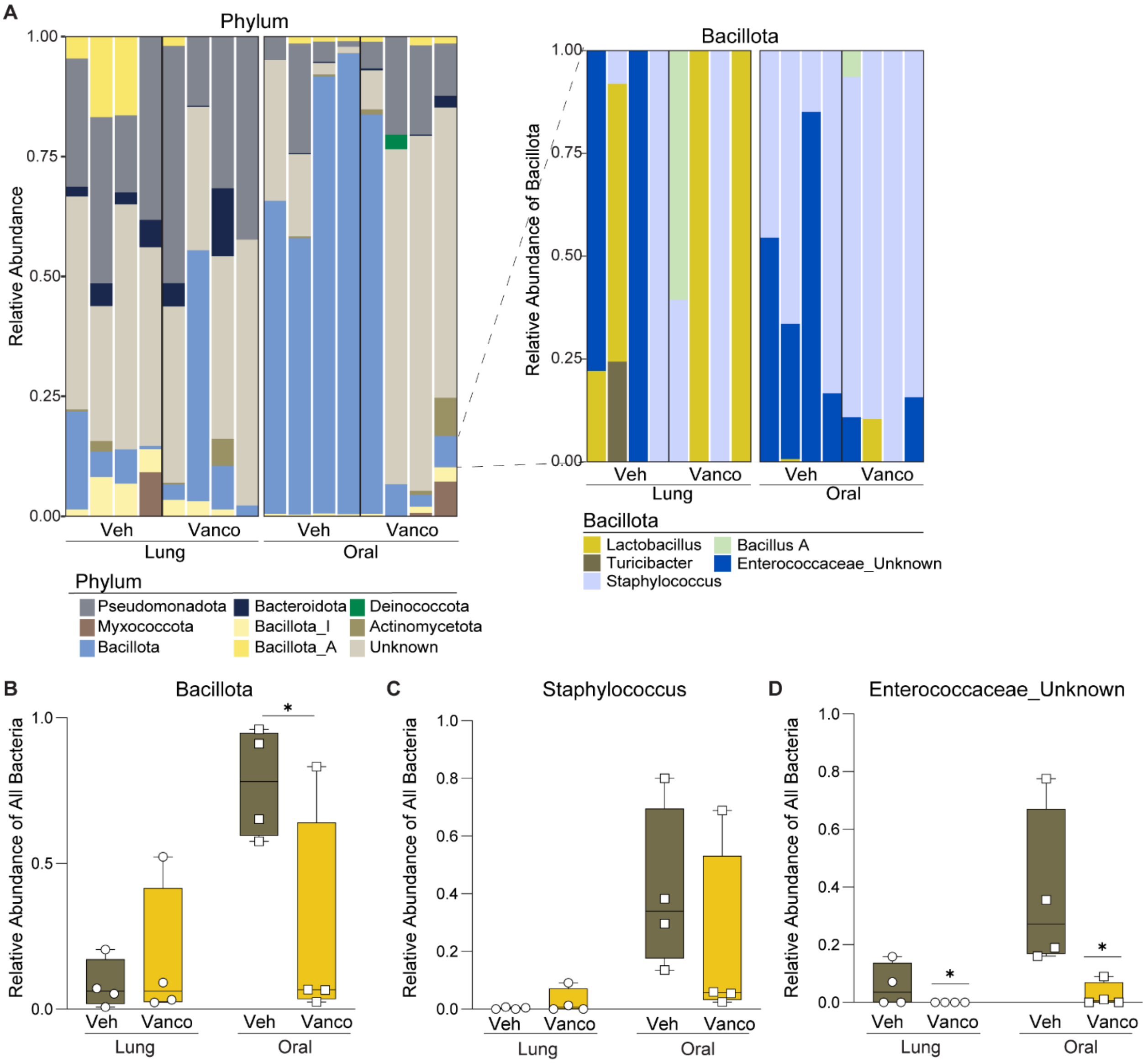


**Supplemental Figure 6. Targeted systemic antibiotics only partially alleviate SCI lung bacterial burden**. (**A**) Stacked bar graphs represent the phyla compositions and a subset stacked bar graph represent different *Bacillota* taxa at the genus level. (**B-D**) Box plots show the relative abundance of different taxa. Each white dot represents one mouse. (**B**) *Bacillota* was significantly reduced by treatment in oral cavity samples (Two-Way ANOVA interaction, $F_{(1,12)} = 5.914$ P= 0.0316, Tukey's multiple comparison test Tongue Veh vs Vanco P=0.0499). No effect of treatment on (**C**) *Staphylococcus,* but a main effect of treatment was seen on (**D**) *Enterococcaceae of an unknown genus* (Two-Way ANOVA interaction, $F_{(1,12)} = 7.384$ P= 0.0187). *16S RNA Sequencing was from a subset of mice in Figure 5, n=4/group). * P<0.05.*
